# Upregulated pexophagy limits the capacity of selective autophagy

**DOI:** 10.1101/2023.05.10.540213

**Authors:** Kyla Germain, Raphaella W. L. So, Joel C. Watts, Robert Bandsma, Peter K. Kim

## Abstract

Selective autophagy is an essential mechanism to maintain organelle integrity and cellular homeostasis through the constant recycling of damaged or superfluous components. While distinct selective autophagy pathways mediate the degradation of diverse cellular substrates including organelles and pathogens, whether these distinct pathways can influence one another remains unknown. We address this question here using pexophagy, the autophagic degradation of peroxisomes, as a model. We demonstrate in cells that upregulated pexophagy exhausts selective autophagy and limits the degradation of both mitochondria and protein aggregates. We confirmed this finding in the pexophagy-mediated form of Zellweger Spectrum Disorder, a rare disease characterized by peroxisome dysfunction. Further, we extend the generalizability of limited selective autophagy by determining that increased aggrephagy reduces pexophagy using a model of Huntington’s Disease. Our findings suggest that the degradative capacity of selective autophagy can become limited by an increased substrate load.

## INTRO

Autophagy is a conserved process responsible for the breakdown of cytoplasmic material, including dysfunctional or superfluous organelles. The canonical autophagy pathway in mammalian cells is macroautophagy (hereafter referred to as autophagy) and is mediated by a vesicular structure called the autophagosome.^1^ Selective autophagy is a form of autophagy wherein a particular cytoplasmic substrate is targeted for degradation by the accumulation of ubiquitin on its outer surface.^2^ Specific autophagy receptor proteins bind to ubiquitinated substrates through a ubiquitin binding domain and help to recruit the autophagy machinery that drives autophagosome formation.^2–4^ Concomitantly, autophagy receptors bind to autophagosomes via an LC3-interacting region, effectively sequestering ubiquitinated substrates within autophagosomes to facilitate their degradation.^2–4^

The selective autophagy pathways that mediate the degradation of various cellular structures have been described in recent years, including the degradation of peroxisomes via pexophagy, mitochondria via mitophagy, and protein aggregates via aggrephagy.^5–7^ However, whether these different selective autophagy pathways influence one another is unknown. Given that distinct selective autophagy pathways require overlapping machinery including autophagosomes and lysosomes for degradation, we postulated that an increase in one selective autophagy pathway may limit the activity of others.

Neurodegenerative disorders marked by a buildup of autophagy substrates including misfolded proteins and damaged organelles may support this hypothesis. Zellweger Spectrum Disorders (ZSD) are a group of a rare, inherited diseases characterized by a loss of peroxisomes leading to a multisystemic disorder that includes neurodegeneration.^8^ Cellularly, the loss of peroxisomes in humans is accompanied by an increase in damaged mitochondria, abnormal lysosomes, and accumulation of lipid droplets, all of which are substrates of autophagy.^9–12^

Studies of three mouse models of ZSD also revealed an accumulation of autophagy substrates including dysfunctional mitochondria and phosphorylated alpha-Synuclein (αS) oligomers in the liver and brain.^13–15^ We have previously demonstrated that *PEX1*, the most commonly mutated gene underlying ZSD, encodes a peroxisomal protein that regulates pexophagy, such that its depletion or mutation leads to upregulated pexophagy and peroxisome loss.^16^ This raises the possibility that increased pexophagy may limit the activity of other selective autophagy pathways in ZSD.

In this study, we examine whether distinct selective autophagy pathways can influence one another and whether the capacity of selective autophagy can become saturated. We find that upregulating pexophagy limits both mitophagy and aggrephagy in an autophagy-dependent manner and may contribute to the pathological protein aggregation and mitochondrial dysfunction in a model of ZSD. Additionally, we provide data to support the generalizability of our findings using a cell model of Huntington’s Disease (HD).

## Results

### Depletion of PEX1 or PEX13 induces pexophagy

To investigate the capacity of selective autophagy, we first generated a cell model of increased pexophagy. The upregulation of pexophagy was used as a case study as it can be genetically induced without significantly disrupting overall cellular homeostasis. It has recently been demonstrated that the loss of PEX1, a component of the peroxisomal AAA-type ATPase, or PEX13, a component of the peroxisomal matrix protein import complex, triggers ubiquitin-dependent pexophagy (Fig 1A).^16, 17^ Knockdown of PEX1 or PEX13 can therefore be used as reliable method to upregulate pexophagy in cultured cells. Expectedly, the depletion of PEX1 or PEX13 in HeLa (Fig. 1B-D) and HEK293 cells (Fig. S1A, B) resulted in 20-40% peroxisome loss, as determined by quantification of the punctate structures visualized by immunofluorescent staining of PMP70, a peroxisome membrane protein. Whereas, depletion of another peroxisome matrix import factor, PEX14, did not affect the number of PMP70 punctate structures in HeLa or HEK293 cells (Fig. 1, S2). The loss of peroxisomes observed in PEX1 or PEX13 depleted cells was autophagy dependent, as co-depletion of the autophagy factor ATG12 prevented peroxisome loss as previously described (Fig 1C, D).^16, 17^

**Figure 1.**
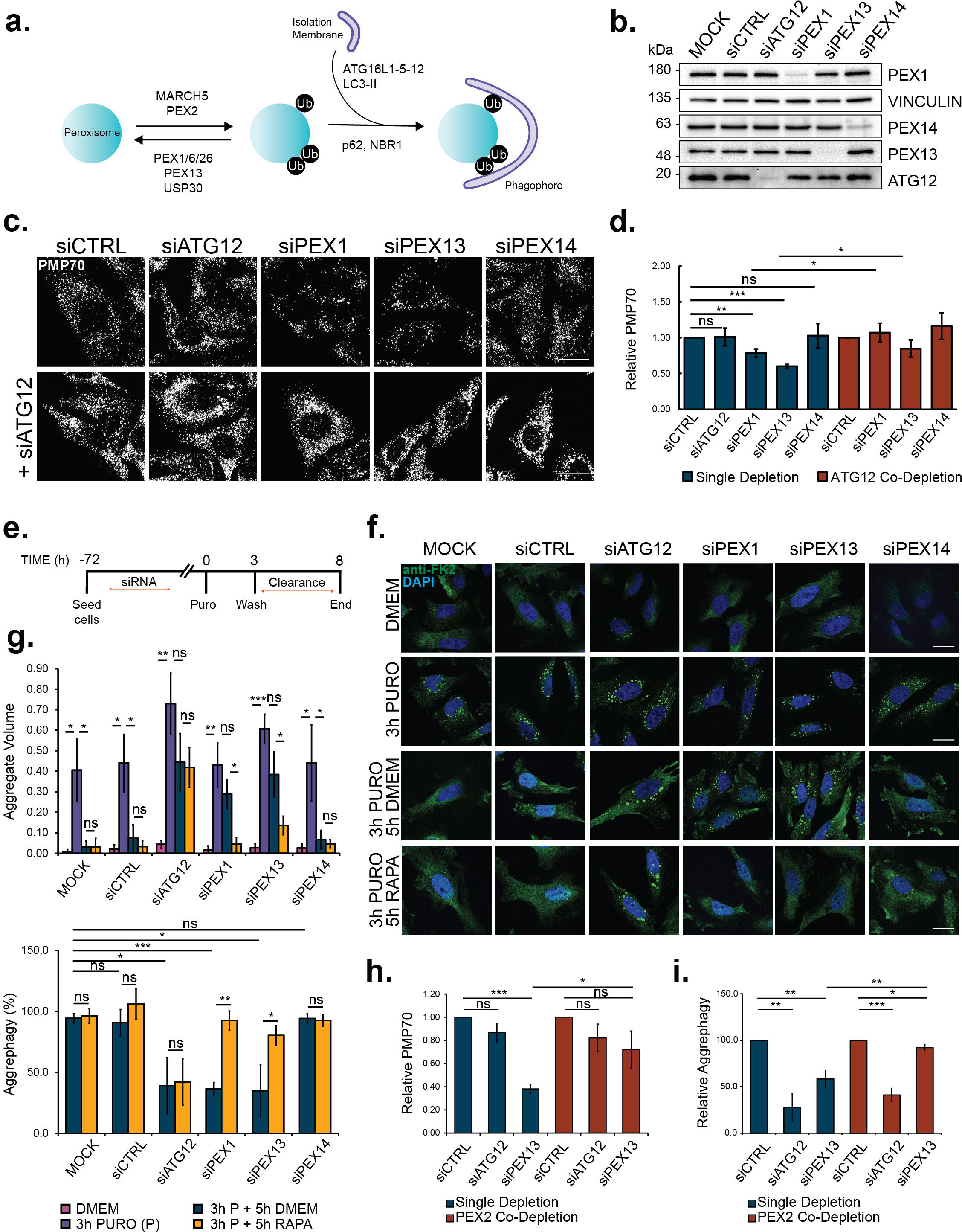
PEX1 or PEX13 depletion impairs puromycin-induced aggrephagy. **(a)** Schematic of mammalian pexophagy. **(b)** Immunoblot of HeLa cells treated with the siRNA and probed for the indicated proteins. **(c)** Representative images of HeLa cells treated with the indicated siRNA and immunostained for the peroxisomal marker, PMP70. Scale bar, 25µm. **(d)** Quantification of PMP70 in (c), relative to control siRNA treated cells. PMP70 was measured by dividing the number of PMP70 puncta by cell volume (see Methods). **(e)** Schematic of aggrephagy assay. HeLa cells were treated with the indicated siRNA prior to aggrephagy induction (see Methods). **(f)** Representative images from cells at each stage in the assay: DMEM, 3-h 5µg mL^-1^ Puromycin, 3-h 5µg mL^-1^ Puromycin followed by 5-h clearance period in DMEM, or 3-h 5µg mL^-1^ Puromycin followed by 5-h clearance period in 2uM Rapamycin. Cells were immunostained with the ubiquitin antibody FK2 and DAPI (blue) Scale bars, 25µm. **(g)** Quantification of aggrephagy assay in (f). Aggregate Volume was calculated by dividing the total volume of FK2 puncta by cell volume. Aggrephagy activity was measured as the % clearance of FK2 aggregates, relative to mock transfected ‘3h P + 5h DMEM’ cells. **(h)** Quantification of relative PMP70 in HeLa cells treated with the indicated siRNAs. **(i)** Aggrephagy activity in HeLa cells treated with the indicated siRNAs, relative to control siRNA-treated condition. Results represent mean from *n*=3 independent trials (30 cells quantified/trial) and error bars represent standard deviation. *P < 0.05, **P <0.01, ***P<0.001; two-tailed unpaired student t-test.

### Increased pexophagy limits aggrephagy

Using our model for upregulated pexophagy, we first asked whether increased pexophagy influences the degradation of protein aggregates via aggrephagy. Specifically, we examined the autophagic clearance of aggresome-like inducible structures (ALIS) from HeLa cells following proteotoxic stress. Aggrephagy was assessed by monitoring the volume of ALIS in cells visualized with an FK2 antibody that binds to poly- and monoubiquitinated proteins, but not free ubiquitin. Proteotoxic stress with 3-h puromycin treatment resulted in a robust production of ALIS (Fig. 1E,F). In control cells transfected with either no siRNA (mock) or non-targeting siRNA, ALIS volume was largely reduced following a 5-h washout/clearance period, as indicated by a significant decline in FK2-aggregate volume (Fig. 1E,F). Whereas, depletion of ATG12 prevented ALIS clearance in line with previous reporting that puromycin-induced ALIS are degraded by autophagy (Fig. 1F, G).^18–20^

To investigate whether increased pexophagy influences aggrephagy, we tested whether PEX1 or PEX13 depletion affected ALIS clearance in HeLa cells. Depletion of PEX1 and PEX13 resulted in decreased aggregate clearance compared to control and PEX14 depleted cells (Fig. 1F,G). We further quantified aggrephagy activity by calculating the percentage clearance of ALIS and observed that similarly to the ATG12 depleted cells, the loss of PEX1 or PEX13 decreased aggrephagy activity compared to control cells (Fig. 1G).

### PEX13 or PEX1 are not required for aggrephagy

Previous work has postulated that peroxisomes, and in particular PEX13, may be required for selective autophagy as the loss of PEX13, but not other peroxisome biogenesis factors PEX14 or PEX19, prevented mitophagy and virophagy in cultured cells and mouse tissue.^21^ To test whether PEX13 is required for aggrephagy, instead of influencing aggrephagy via increased pexophagy, we co-depleted HeLa cells of the peroxisomal E3 ubiquitin ligase that is required for the induction of pexophagy, PEX2 (Fig. 1A).^17, 22^ We corroborated previous reporting that co-depletion of PEX13 and PEX2 rescues the peroxisome loss observed in cells depleted of PEX13 alone, indicating that PEX2 co-depletion is sufficient to inactivate pexophagy in PEX13-depeleted cells (Fig. 1H).^17^ Co-depletion of PEX2 with PEX13 also rescued aggrephagy activity compared to cells depleted of PEX13 alone, supporting that the upregulation of pexophagy and not the loss of PEX13 function impaired aggrephagy (Fig. 1I).

To further validate that upregulated pexophagy is responsible for the retarded aggrephagy activity, we induced pexophagy without depleting any peroxisomal genes. It was previously shown that targeting a ubiquitin motif to the cytosolic face of the peroxisomal membrane by ectopically expressing PMP34-GFP-UBKo can induce pexophagy.^23^ PMP34-GFP-UBKo is a chimera consisting of the peroxisomal membrane protein, PMP34, fused to a GFP and a ubiquitin moiety whose lysine residues were mutated to arginine to prevent polyubiquitination.^23^ As previously shown^23^, expression of PMP34-GFP-UBKo resulted in significant peroxisome loss compared to cells expressing the non-ubiquitin tagged PMP34-GFP (Fig. 2A,B). When subjected to the puromycin-ALIS aggrephagy assay, cells expressing PMP34-GFP-UBKo had less aggrephagy activity compared to PMP34-GFP expressing cells (Fig. 2C-E), similar to the results observed in PEX1 and PEX13-depleted cells (Fig. 1).

**Figure 2.**
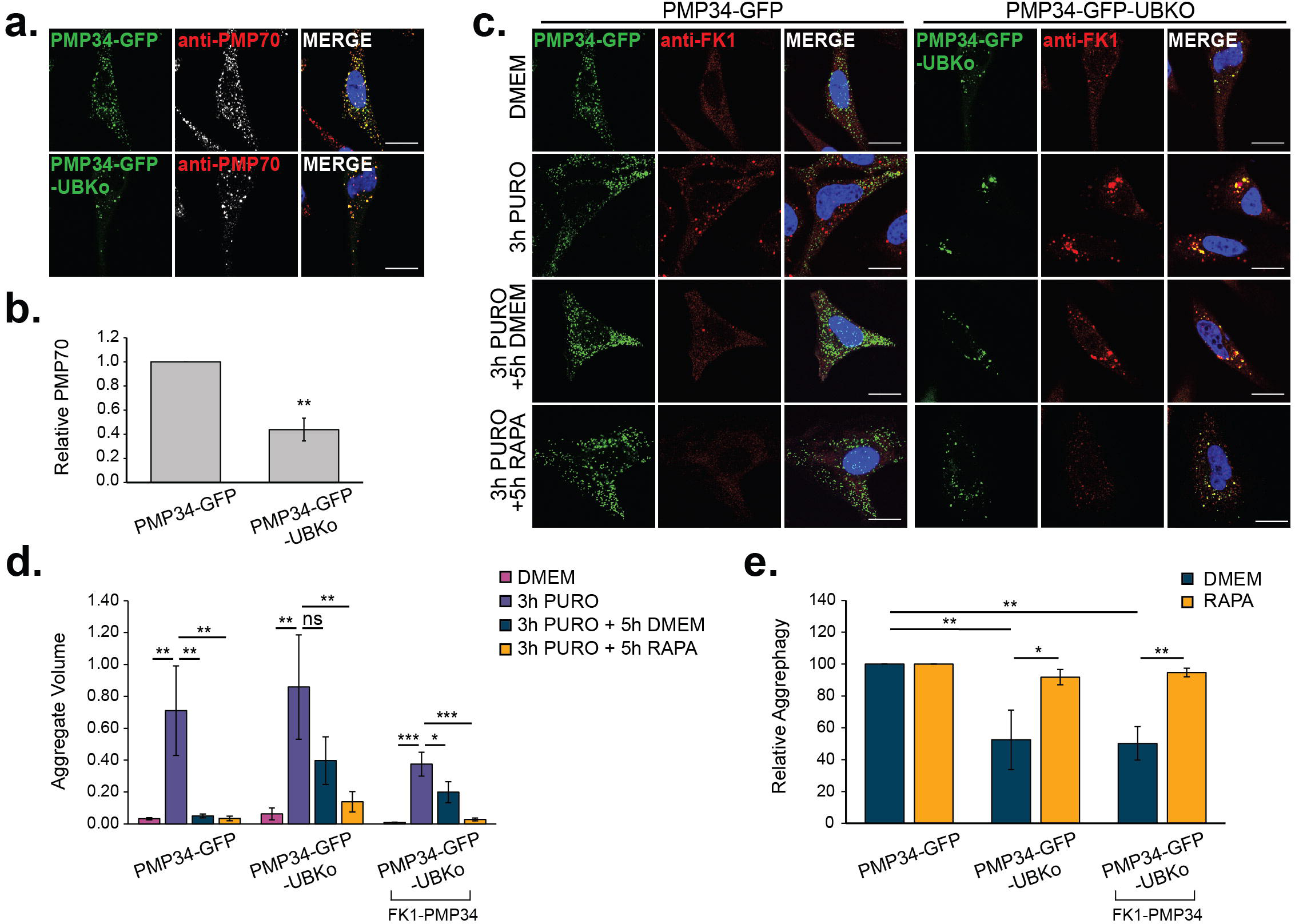
PMP34-GFP-UBKo expression impairs puromycin-induced aggrephagy. **(a)** HeLa cells transfected with either PMP34-GFP or PMP34-GFP-UBKo 24-h before fixation and immunostaining for PMP70. Scale bars, 20µm. **(b)** Quantifications of PMP70 in (a), relative to PMP34-GFP condition. PMP70 was calculated by dividing the number of PMP70 puncta by cell volume (see Methods). **(c)** HeLa cells transfected with either PMP34-GFP or PMP34-GFP-UBKo 24-h before aggrephagy induction. Representative images from cells at each stage in the assay: DMEM, 3-h 5µg mL^-1^ Puromycin, 3-h 5µg mL^-1^ Puromycin followed by 5-h clearance period in DMEM, or 3-h 5µg mL^-1^ Puromycin followed by 5-h clearance period in 2uM Rapamycin DMEM. Cells were immunostained for FK1. Scale bars, 25µm. **(d)** Quantification of Relative Aggregate Volume in (c). Relative Aggregate Volume was calculated by dividing the total volume of FK1 puncta by cell volume. **(e)** Quantification of aggrephagy activity in (c), relative to PMP34-GFP. Aggrephagy activity was measured as the % clearance of FK1 aggregates. Results represent mean from *n*=3 independent trials (30 cells quantified/trial) and error bars represent standard deviation. *P < 0.05, **P <0.01; two-tailed unpaired student t-test.

Surprisingly, we observed colocalization of PMP34-GFP-UBKo and ALIS following proteotoxic stress, suggesting that peroxisomes and ALIS may be clustered to allow for autophagy. As ALIS were here visualized with an FK1 antibody that binds to only poly-, and not monoubiquitinated proteins, labelling of the monoubiquitinated PMP34-GFP-UBKo should be subverted. However, to ensure that our quantification of FK1 represented true ALIS and not ubiquitinated peroxisomes, we subtracted the volume of PMP34-GFP-UBKo structures from the volume of FK1-labelled structures per cell (Fig. 2D). When adjusted for the PMP34-GFP-UBKo structures, we still observed less aggrephagy activity in cells expressing PMP34-GFP-UBKo compared to those expressing PMP34-GFP (Fig. 2E)

Finally, we tested whether the loss of functional peroxisomes may be responsible for limiting aggrephagy by measuring ALIS clearance in PEX19 deficient peroxisome biogenesis disorder (PBD) 399-T1 human fibroblast cells. PEX19 is an essential peroxisome biogenesis factor, wherein its absence peroxisome biogenesis is abolished.^24^ We confirmed that PBD399-T1 cells lack peroxisomes by immunoblot analysis of PMP70 (Fig. S2A,B). While PBD399-T1 cells lack peroxisomes they still express some peroxin proteins, therefore we treated the cells with siRNA to ensure a loss of PEX1 and PEX13 (Fig. S2A). Despite lacking peroxisomes, PBD399-T1 cells were able to clear puromycin-induced ALIS, supporting their capacity for aggrephagy, while disrupting autophagy by ATG12 depletion prevented ALIS clearance (Fig. S2C-E).

Collectively, these data indicate that upregulation of pexophagy and not the loss of PEX1, PEX13 or peroxisome function elicit impaired aggrephagy.

### Upregulating autophagosome formation rescues aggrephagy in pexophagy-induced cells

Limited aggrephagy in conditions of increased pexophagy suggest that an upsurge of pexophagy may be sequestering the available autophagosome initiation/biogenesis machinery, thus limiting aggrephagy from occurring effectively. To test this hypothesis, we repeated our aggrephagy assays in the presence of the mTORC1 inhibitor, rapamycin, to upregulate autophagosome formation and autophagy flux. We confirmed that 5-h of rapamycin treatment robustly inhibited mTORC1 activity in PEX1 or PEX13 cells by measuring phosphorylation of its downstream target, p70 S6 Kinase (S6K) (Fig. S3).

When cells were supplemented with rapamycin during the 5-h clearance period of the ALIS assay, we observed a statistically significant improvement in aggrephagy activity in both PEX1 and PEX13, but not ATG12 depleted cells (Fig. 1E-G). Akin to the PEX1 and PEX13-depleted cells, rapamycin treatment improved the clearance of ALIS from PMP34-GFP-UBKo-expressing cells (Fig. 2C-E). Taken together, these data indicate that increased pexophagy limits aggrephagy in an autophagy-dependent manner.

### Increased pexophagy limits mitophagy

We next addressed whether increased pexophagy influences mitophagy. Specifically, we examined the effects of increased pexophagy on Parkin-mediated mitophagy induced with the mitochondrial respiration inhibitors, oligomycin and antimycin A1 (OA)^25^, in HEK293 cells stably expressing GFP-Parkin. In mock control or non-targeting siRNA-treated cells, mitochondrial clearance was observed following 8-h of OA treatment, as assessed by immunoblot of outer mitochondrial membrane (OMM; MFN2), inner mitochondrial membrane (IMM; CVα) and matrix proteins (Hsp60) (Fig. 3A-D). Inhibition of autolysosome fusion and degradation with bafilomycin A1 (BafA1) did not prevent the degradation of OMM proteins, supporting their proteasomal degradation^5^, but did prevent the degradation of IMM and matrix proteins, confirming their autophagic degradation (Fig. 3A-D). We further observed mitophagy following 8-h OA treatment with immunofluorescent imaging and intensity quantifications of CVα (Fig. 3E,F).

**Figure 3.**
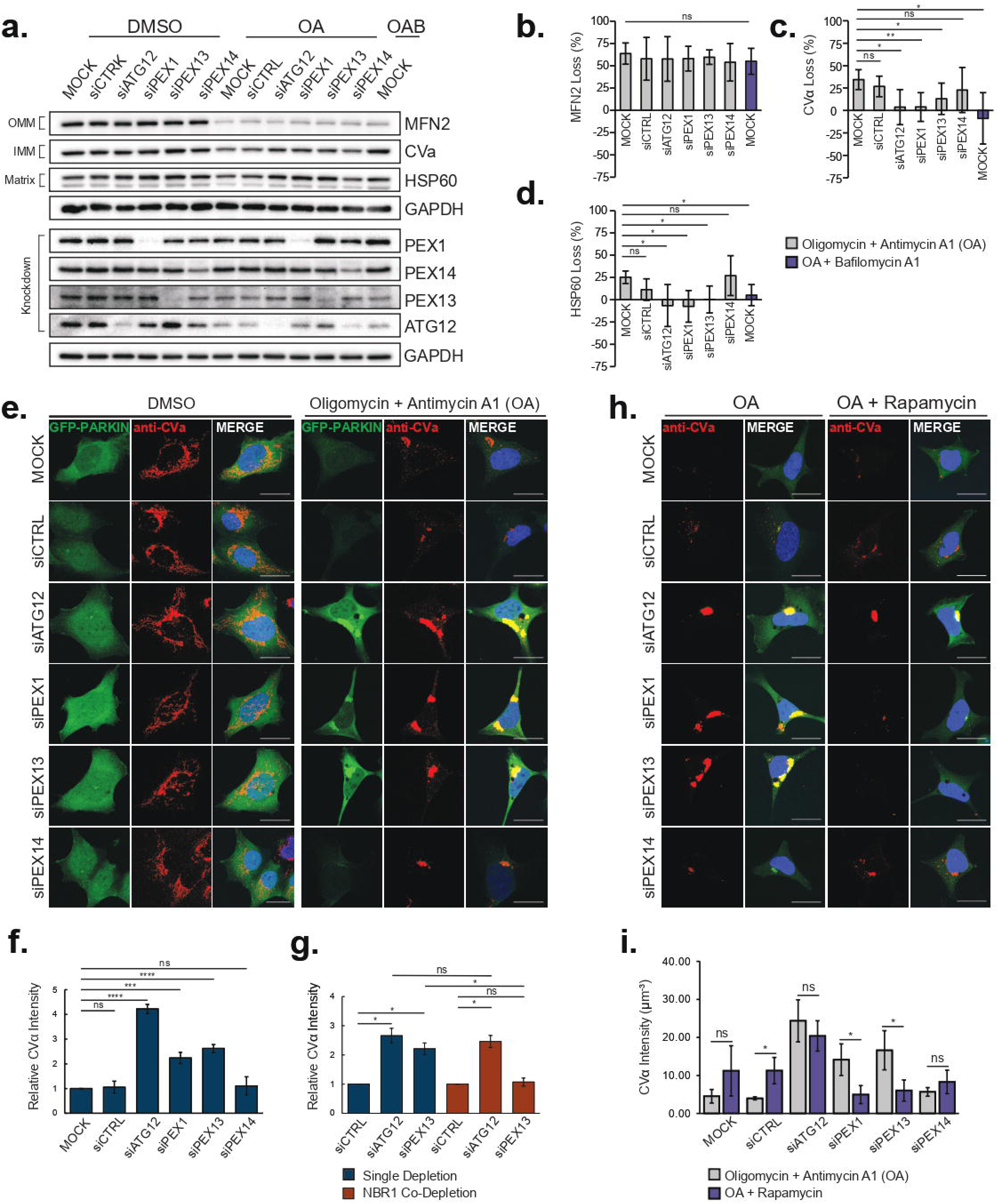
PEX1 or PEX13 depletion impairs mitophagy. **(a)** Immunoblots of GFP-Parkin HEK293 cells subjected to siRNA knockdown prior to treatment for 8-h with DMSO, 2.5µM Oligomycin and 250nM Antimycin A1 (OA), or OA + 250nM Bafilomycin A1. Immunoblots were probed for outer mitochondrial membrane marker MFN2, inner mitochondrial membrane marker CVα, mitochondrial matrix protein HSP60, and siRNA targets: PEX1, PEX13, PEX14, and ATG12. **(b-d)** Percentage loss densitometry quantification of the indicated bands normalized to loading control GAPDH. Bands were quantified using ImageJ software from *n=4* independent trials. **(e)** Representative confocal fluorescent images of GFP-Parkin HEK293 cells treated as in (a). Cells immunostaining for CVα as indicated. Blue= DAPI; Scale bars, 25µm. **(f)** Quantification of CVα after OA treatment in (e), relative to mock transfected cells. CVα was quantified by dividing the total CVα florescence intensity by cell volume. **(g)** Quantification of CVα after 8-h OA treatment in GFP-Parkin HEK293 cells treated with the indicated siRNA (representative images in Fig. S3). CVα is depicted relative to control siRNA treated cells for single depletions, or control + NBR1 siRNA for co-depletions. **(h)** Representative images of GFP-Parkin HEK293 cells treated with the indicated siRNA prior to treatment for 8-h with either OA, or OA + 2µM Rapamycin. Cells were immunostained for mitochondria marker, CVα. Scale bars, 25µm. **(i)** Quantification of CVα in (h). Results represent mean from *n*=3 independent trials (30 cells quantified/trial) and error bars represent standard deviation. *P < 0.05, ***P <0.001, ****P<0.0001; two-tailed unpaired student t-test.

To further validate that the loss of mitochondrial proteins observed following OA treatment was due to mitophagy, we depleted cells of ATG12 to prevent autophagy. The loss of ATG12 decreased mitochondria clearance following 8-h mitophagy induction, as evident from the reduced clearance of IMM and matrix proteins and the four-fold higher fluorescent intensity of CVα in cells depleted of ATG12 compared to controls (Fig. 3A-F). When we induced pexophagy by depletion of PEX1 or PEX13 before the activation of mitophagy, we also observed impaired mitochondrial clearance compared to control cells (Fig. 3A-F). However, this impairment was not observed in cells depleted of PEX14, where pexophagy is not upregulated (Fig. 3A-F). The reduced mitophagy observed with ATG12, PEX1, or PEX13 depletion was not due to defective Parkin recruitment, as Parkin colocalized with mitochondria in OA-treated cells (Fig. 3E) and MFN2, an early substrate of Parkin, was cleared at comparable levels to control cells (Fig. 3A-D). However, the stabilization and colocalization of GFP-Parkin with CVα upon OA treatment in ATG12, PEX1, and PEX13 suggested a defect in the autophagic turnover of mitochondria.

To confirm that the observed mitophagy impairment was a direct result of upregulated pexophagy and not due to the loss of PEX13 function, we repeated our assay in cells with inactivated pexophagy. We prevented peroxisome loss by co-depleting cells of PEX13 and NBR1, an autophagy receptor required for the sequestration of ubiquitinated peroxisomes, but not mitochondria, within autophagosomes.^25, 26^ Cells co-depleted of PEX13 and NBR1 therefore lack pexophagy and peroxisome loss, as previously shown.^17^ Co-depletion of PEX13 and NBR1 prevented the defect in mitochondrial clearance compared to depletion of PEX13 alone (Fig. 3G, Fig. S1C,D), supporting that the defect in mitophagy is a result of upregulated pexophagy and not the loss of PEX13 function.

We next asked whether rapamycin treatment could improve mitochondrial clearance in conditions of upregulated pexophagy. Rapamycin treatment improved the clearance of mitochondria in cells depleted of PEX1 and PEX13, but not in ATG12 depleted cells, as indicated by CVα fluorescence intensity (Fig. 3H,I), further illustrating that upregulated pexophagy retards mitophagy.

### Increased pexophagy impairs the clearance of protein aggregates in ZSD models

Aggrephagy is required for the removal of pathogenic protein aggregates such as αS, the protein commonly known for its oligomerization and deposition within pathological inclusions in Parkinson’s Disease (PD).^27^ αS oligomerization and deposition within cytoplasmic inclusions has been described in a number of ZSD mouse models including PEX13 deficient mice.^13^ Therefore, we next asked whether upregulated pexophagy contributes to the accumulation of αS in ZSD by testing whether PEX1 or PEX13 depletion impairs the clearance of αS pre-formed fibrils (PFF) *in vitro*. αS PFF transfected into HeLa cells formed large aggregates within 4-h and were cleared within 8-h after washing out the transfection complex (clearance period) (Fig 4B: α-Syn PFF vs α-Syn+8h DMEM). Inhibiting lysosomal protease activity by the addition of Leupeptin and E-64 during the clearance period resulted in the persistence of αS aggregates that colocalized with LC3, supporting their autophagic degradation (Fig. 4B).

**Figure 4.**
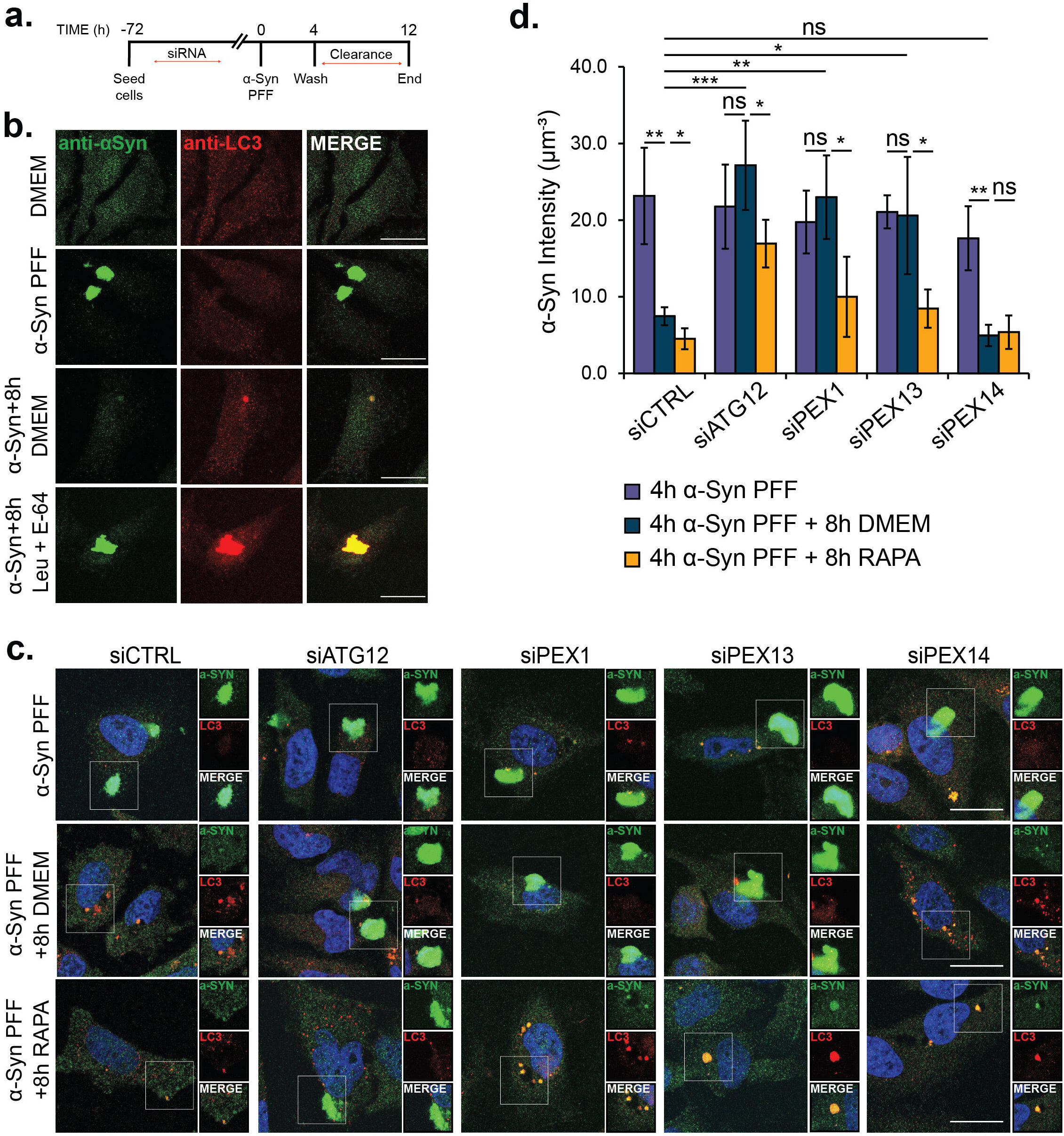
PEX1 or PEX13 depletion impairs the autophagic clearance of alpha-Synuclein fibrils. **(a)** Schematic of alpha-Synuclein assay. 72-h after siRNA transfection: 4-h transfection with alpha-Synuclein pre-formed fibrils (αS _PFF_); 4-h αS PFF followed by an 8-h clearance period in either DMEM or 0.25mM Leupeptin and 2uM E-64. **(b)** Immunofluorescent images of HeLa cells at each stage in the assay in (a): Cells were immunostained for αS and LC3. Scale bars, 25µm. **(c)** HeLa cells treated with the indicated siRNA before αS PFF transfection. Representative images from cells transfected for 4-h with αS PFF, 4-h αS PFF followed by an 8-h clearance period in either DMEM or 2uM Rapamycin. Cells were immunostained for αS and LC3. Scale bars, 25µm. **(d)** Quantification of relative αS intensity in (c), calculated by dividing the total αS florescence intensity by cell volume. Results represent the mean from *n*=4 independent trials, and error bars represent standard deviation (30 cells quantified/trial). *P < 0.05, **P <0.01, ***P<0.001; two-tailed unpaired student t-test.

We observed that cells depleted of ATG12, PEX1, or PEX13 displayed impaired clearance of αS compared to control or PEX14 depleted cells, manifesting as significantly higher αS fluorescence intensity (Fig. 4A,C,D). Notably, αS did not colocalize with LC3 as observed by immunofluorescent imaging (Fig. 4C). However, treatment with rapamycin improved αS clearance in PEX1 and PEX13 depleted with a reduction in αS fluorescence intensity, as well as promoted colocalization of αS with LC3 (Fig. 4C,D). These results indicate that increased pexophagy impairs the clearance of αS by aggrephagy.

To evaluate whether aggrephagy was also impaired in a ZSD model with increased pexophagy, we repeated the puromycin-induced ALIS assay in PEX1^G843D^ patient fibroblast cells. PEX1^G843D^ is the most common mutation underlying ZSD; a null mutation resulting in peroxisome loss through increased pexophagy.^16, 28, 29^ Compared to control human fibroblast cells, PEX1^G843D^ cells exhibited a reduction in aggrephagy activity, which was improved with rapamycin supplementation (Fig. 5). As a control, we also measured aggrephagy activity in another ZSD patient fibroblast cell line harboring a nonsense mutation in the critical peroxisome biogenesis gene, PEX3-R53ter. PEX3-R53ter cells therefore lack peroxisomes because of defective peroxisome biogenesis.^30^ We observed aggrephagy levels in the PEX3-R53ter fibroblasts similar to control fibroblasts, supporting that the aggrephagy defects in the PEX1^G843D^ cells are due to upregulated pexophagy, not a lack of peroxisomes.

**Figure 5.**
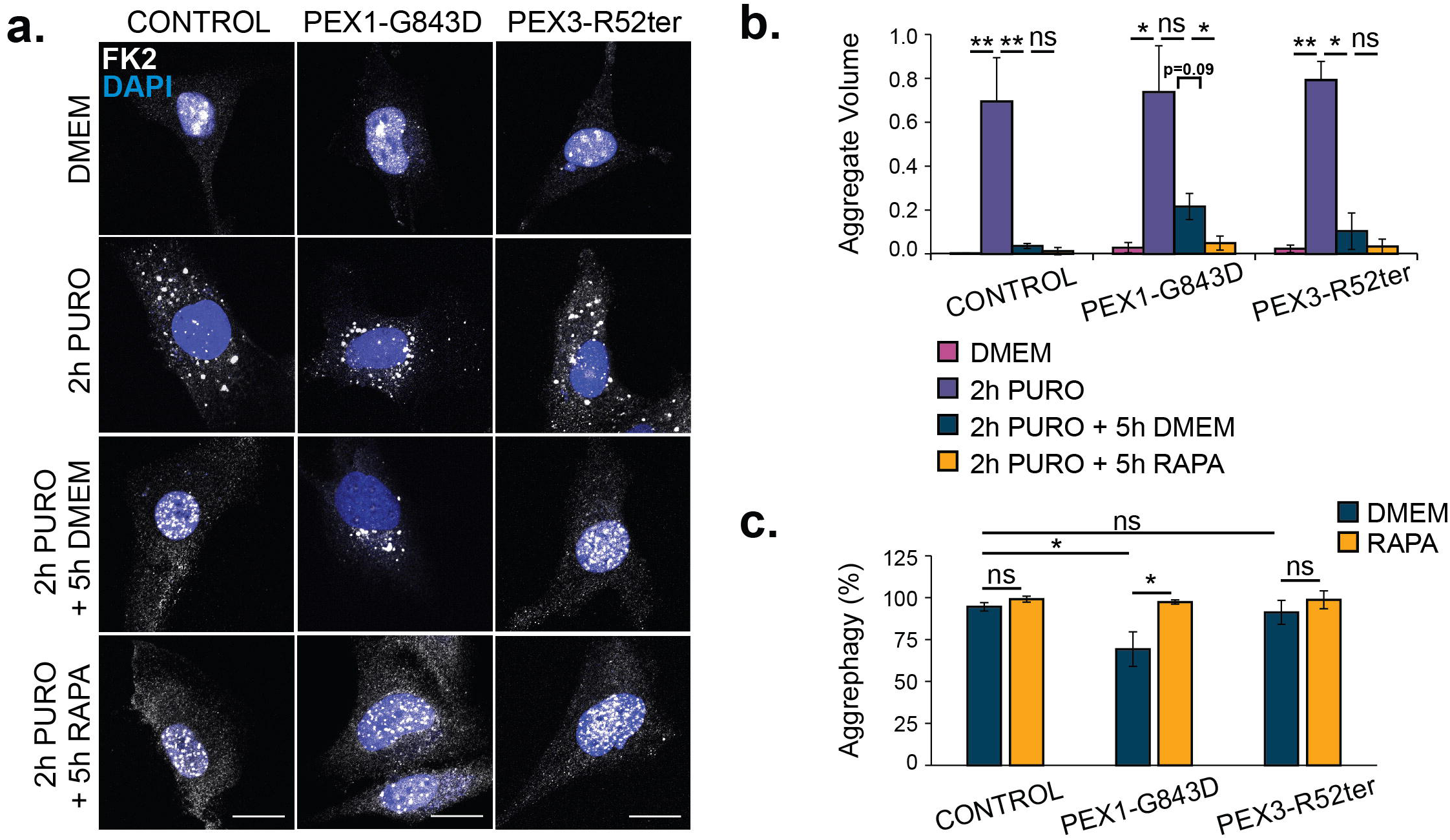
Zellweger Spectrum Disorder patient fibroblasts exhibit impaired puromycin-induced aggrephagy. **(a)** Representative images of control, PEX1^G843D^, or PEX3-R52Ter ZSD fibroblasts at each stage in the aggrephagy assay. Cells were immunostained with the antibody FK2. Scale bars, 25µm. **(b)** Quantification of Relative Aggregate Volume in (a). Relative Aggregate Volume was calculated by dividing the total volume of FK2 puncta by cell volume. **(c)** Quantification of aggrephagy activity in (a), relative to control fibroblasts. Aggrephagy activity was measured as the % clearance of FK2 aggregates. Results represent mean from *n*=3 independent trials (30 cells quantified/trial) and error bars represent standard deviation. *P <0.05; two-tailed unpaired student t-test.

### mTORC1 is not activated by PEX1 or PEX13 loss

The ability of rapamycin to rescue mitophagy and aggrephagy in cells with upregulated pexophagy suggests that an mTORC1-regulated step of autophagy is limited. Rapamycin treatment relieves the mTORC1-mediated inhibition of autophagy, resulting in the activation of autophagosome and lysosome biogenesis and the upregulation of certain autophagy receptors.^31, 32^ Therefore, we next investigated the specific mTORC1-regulated selective autophagy machineries that may be limited during upregulated pexophagy.

First, we investigated whether the depletion of either PEX1 or PEX13 resulted in hyperactive mTORC1 thus further restricting autophagy, as has been shown to limit selective autophagy in some lysosomal storage disorders.^33^ However, we observed no significant difference in the phosphorylation state of mTORC1 and its downstream substrate p70-S6K between the various siRNA treatments (Fig. S3).

### Autophagy receptors are not uniformly limited during upregulated pexophagy

Both protein aggregates and peroxisomes are targeted for autophagy by the receptors NBR1 and p62, whereas mitophagy primarily requires OPTN.^5, 7, 26^ While p62 and NBR1 help to cluster ubiquitinated mitochondria, they are not required for mitophagy.^5^ Therefore, we asked whether increased pexophagy delays the degradation of other substrates by consuming autophagy receptors. To test this, we measured the abundance of NBR1, p62, and OPTN during upregulated pexophagy to observe whether their abundance was diminished. In cells depleted of ATG12, we observed increased abundance of all autophagy receptors compared to mock-transfected cells, confirming impaired autophagic flux (Fig. 6A-D). In cells depleted of either PEX1 or PEX13, we observed that OPTN abundance was increased to similar levels as seen with ATG12 depletion, supporting a defect in mitophagy flux but not a limitation in receptor availability (Fig. 6D-G). As previously shown, the depletion of PEX1 resulted in decreased p62 and NBR1 abundance, which could suggest that these receptors become limiting during PEX1 depletion (Fig. 6D-G).^16^ However, PEX13 depletion caused no change to NBR1 abundance and an increase in p62 abundance (Fig. 6D-G). Since cells depleted of PEX14 exhibited a similar increase in p62 as those depleted of PEX13, it is possible that the upregulation of p62 is not related to pexophagy but to other functions shared by PEX13 and PEX14 which together form the peroxisome import pore.^34^ Taken together, these findings demonstrate that autophagy receptor abundance is affected in PEX1 or PEX13 depleted cells, however their changes do not uniformly support that autophagy receptors are a rate-limiting machinery for selective autophagy.

**Figure 6.**
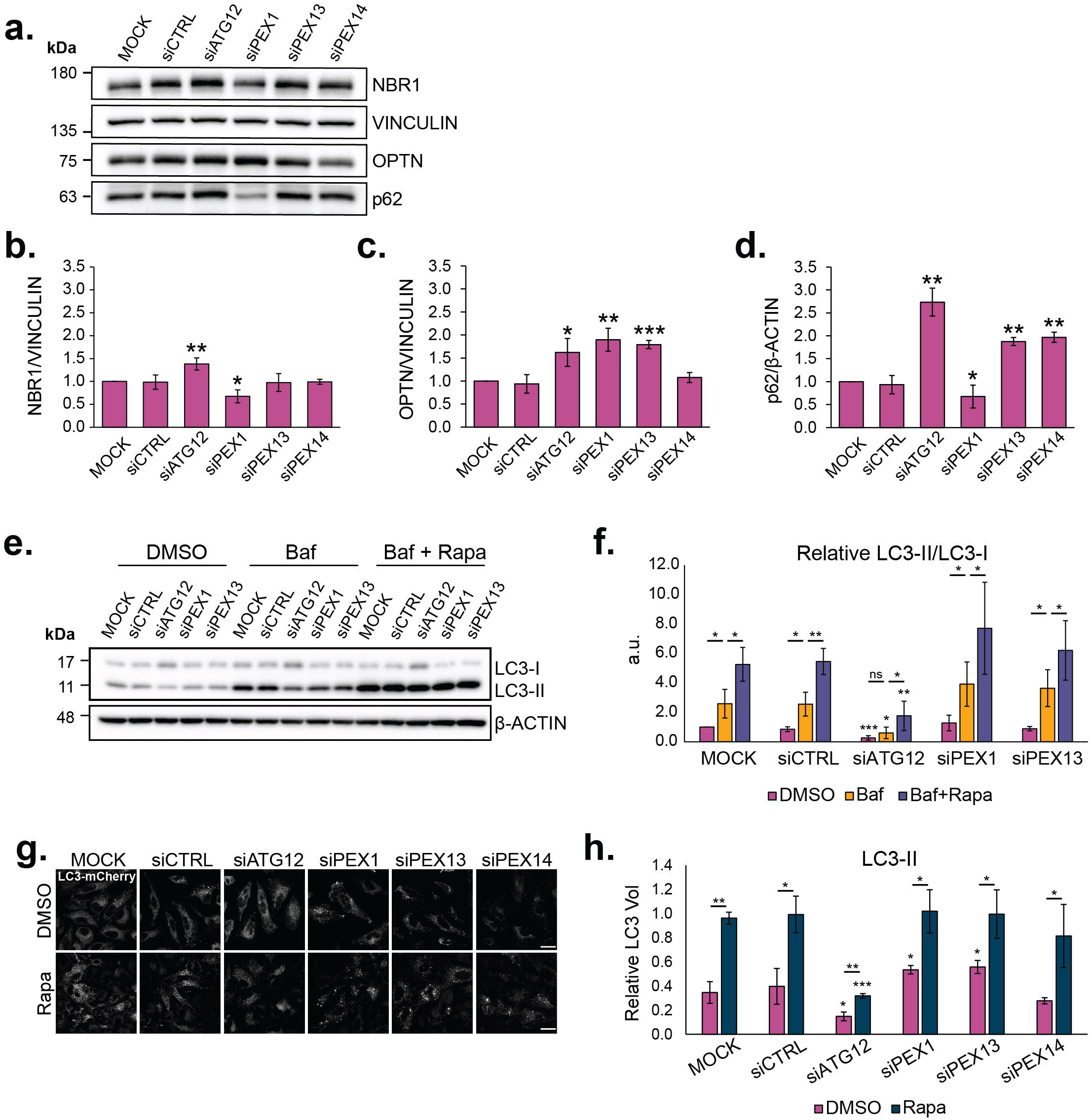
Rapamycin treatment induces autophagy flux in PEX1 and PEX13 depleted cells. **(a)** Immunoblot of HeLa cells treated with the indicated siRNA and probed for NBR1, OPTN, p62 and Vinculin. **(b-d)** Densitometry quantification of (b) NBR1, (c) OPTN, and (d) p62 normalized to Vinculin and relative to MOCK. **(e)** Immunoblot of HeLa cells treated with the indicted siRNA and grown in either DMSO, 200nm Bafilomycin, or 2µM Rapamycin + 200nM Bafilomycin A1 for 5-h. **(f)** Densitometry quantification of LC3-II/LC3-I ratio in (e), relative to MOCK DMSO condition. **(g)** Representative images of HeLa cells treated with the indicated siRNA, transfected with LC3-mCherry, and grown in either DMSO or 2µM Rapamycin for 5-h. Cells were digitonin permeabilized prior to fixation. Scale bars, 25µm. **(h)** Quantification of the volume of LC3 puncta, relative to cell volume in (j). All results represent mean from *n*=3 independent trials and error bars represent standard deviation. *P < 0.05, **P <0.01, ***P<0.001; two-tailed unpaired student t-test.

### Autophagosome formation is not suppressed in PEX1 or PEX13 depleted cells

Next, we asked whether autophagosome formation or flux may be suppressed in PEX1 or PEX13 depleted cells. We measured basal autophagy flux by treating cells with BafA1 and observed no significant changes in the LC3-II:I ratio between PEX1 or PEX13 depleted cells and control cells (Fig. 6E,F). Instead, we observed an increasing trend in the LC3-II:I ratio which could suggest that PEX1 and PEX13 depleted cells undergo increased autophagy flux (Fig. 6E,F). Rapamycin treatment significantly increased the LC3-II:I ratio, demonstrating that PEX1 and PEX13 depleted cells have the capacity to upregulate autophagosome biogenesis (Fig. 6E,F).

We confirmed these findings by monitoring the abundance of LC3 puncta by immunofluorescence in DMSO or rapamycin treated cells. Cells were transfected with LC3- Cherry and permeabilized with digitonin prior to fixation in order to wash out cytosolic LC3-I, such that only membrane bound LC3-II remains (Fig. 6G). Here we observed that PEX1 or PEX13 depleted conditions had an increase in LC3-II puncta compared to mock-treated cells, supporting increased basal autophagosomes formation in these conditions (Fig. 6G,H). Further, rapamycin treatment resulted in a large expansion of LC3-II puncta in all cells, again demonstrating their capacity to upregulate autophagosomes (Fig. 6I,J). Collectively, these findings suggest that PEX1 and PEX13 depletion does not inhibit autophagosome formation or flux, but instead results in a modest increase in basal autophagosome levels.

### Lysosomes are not defective during upregulated pexophagy

Next, we investigated whether lysosome availability may limit selective autophagy during conditions of upregulated pexophagy. One of the final processes of autophagy is the regeneration of spent lysosomes.^35^ To visualize and quantify mature lysosomes, we compared the number and colocalization of LAMP1A positive structures with the acidic lysosome marker cresyl violet, an acidotropic fluorescent marker that is resistant to photobleaching and photoconversion.^36^ As previously shown, neutralizing the lysosome with Concanamycin (CCa) and NH_4_Cl abolished cresyl violet structures but not LAMP1A structures (Fig. 7A-C).^36^ However, the induction of autophagy with either rapamycin or torin1 for 24-h resulted in an increase in cresyl violet structures compared to untreated cells (Fig. 7A-C). We observed an increase in LAMP1A structures in Torin1 treated, but not rapamycin treated cells compared to untreated cells (Fig. 7A-C). This supports previous reporting that Torin1 and not rapamycin prevents the mTORC1-mediated inhibition of TFEB, the lysosome biogenesis transcription factor, to promote lysosomal biogenesis.^37^ Further, quantification of LAMP1A and cresyl violet colocalization revealed increased colocalization in rapamycin and torin1 treated cells, and decreased colocalization in CCa + NH_4_Cl treated cells (Fig. 6D).

**Figure 7.**
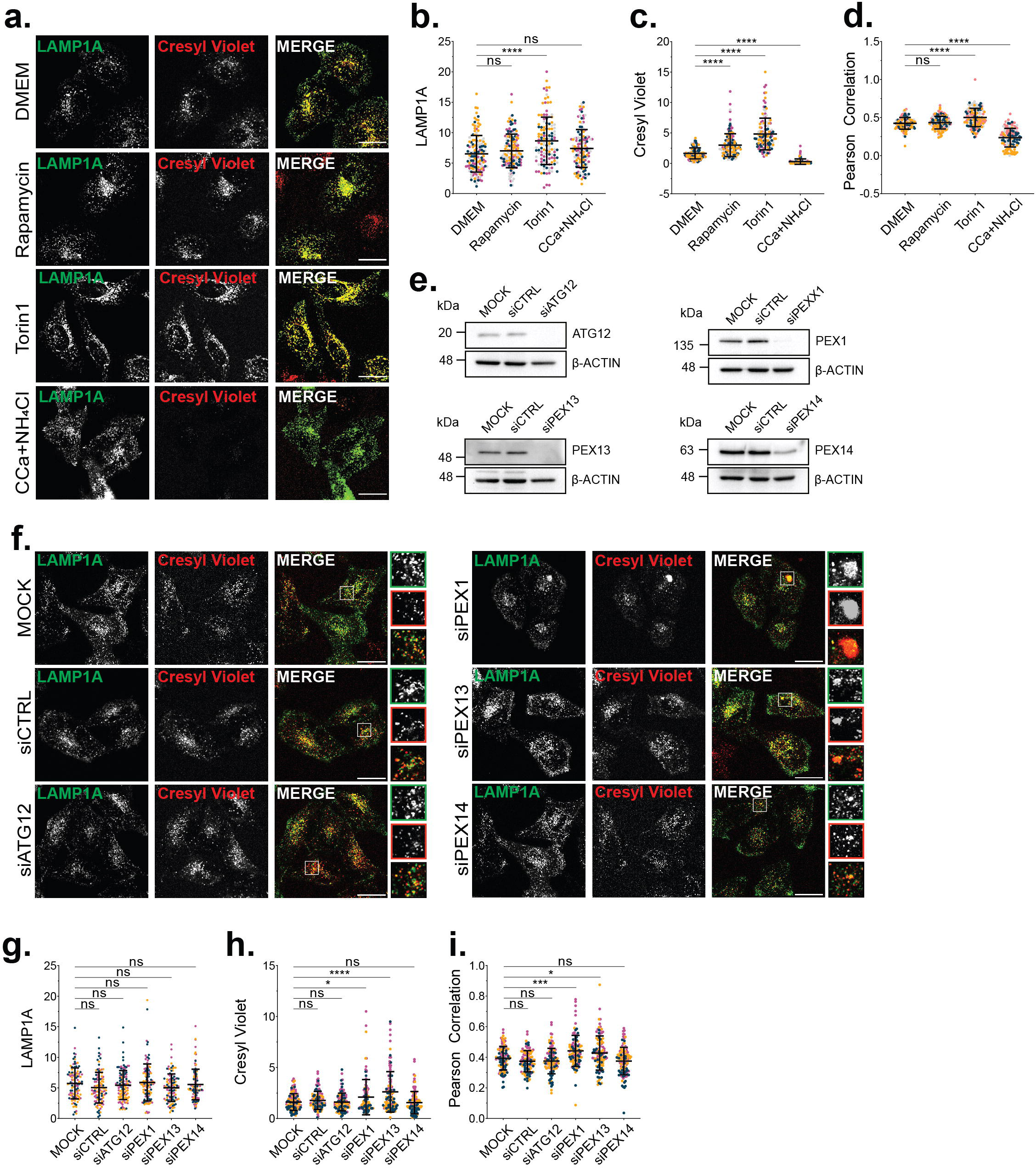
PEX1 or PEX13 depletion results in an expansion of acidic lysosomes. **(a)** Immunofluorescent images of HeLa cells electroporated with LAMP1A and grown in DMEM for 24-h. Cells were treated with either 100nM Rapamycin or Torin for 24-h, or Concanamycin and NH_4_Cl for 1-m, and subjected to Cresyl Violet treatment for 5-m prior to live imaging (see Methods). Scale bar, 25µm. **(b-d)** Quantification of (a) for LAMP1A (b) and Cresyl Violet (c) relative to cell volume, and Pearson Correlation coefficient of LAMP1A and Cresyl Violet (d). Each point denotes one cell, and each color represents one trial of 4 independent trials (30 cells quantified/trial). **(e)** Immunoblot of HeLa cells electroporated with the indicated siRNA and probed for β-Actin and either ATG12, PEX1, PEX13, or PEX14. **(f)** Immunofluorescent images of HeLa cells electroporated with LAMP1A and the indicated siRNA and grown in DMEM for 24-h, and labelled with Cresyl Violet prior to live imaging (see Methods). Scale bar, 25µm. Quantification of **(g)** LAMP1A or **(h)** Cresyl Violet relative to cell volume, and **(i)** Pearson Correlation coefficient of LAMP1A and Cresyl Violet in (f) (see Methods). Each point denotes one cell, and each color represents one trial from *n*=4 independent trials (30 cells quantified/trial). *P < 0.05, **P <0.01; One-Way ANOVA.

Using this assay, we addressed whether upregulated pexophagy affects the population of mature lysosomes. We did not observe a decrease in mature lysosomes in either PEX1 or PEX13 depleted cells (Fig. 6F-I). Instead, we observed an increase in cresyl violet positive structures which was not observed in ATG12 or PEX14 depleted cells (Fig. 6F-H). Moreover, cells depleted of PEX1 or PEX13 had an increase in LAMP1A and cresyl violet colocalization compared to mock and control siRNA treated cells, supporting an expansion of mature lysosomes with PEX1 or PEX13 depletion (Fig. 6I). Morphologically, some PEX1 and PEX13 depleted cells contained large, perinuclear structures that were positive for LAMP1A and cresyl violet, however these structures were not consistently present in cells (Fig. 6F). Together, these data suggest that lysosome availability is not limiting during upregulated pexophagy.

Collectively, these studies suggest that upregulated pexophagy does not inhibit selective autophagy of other substrates by mTORC1 hyperactivity, consumption of autophagy receptors or reducing lysosomal availability. Instead, it suggests that the increased usage of available autophagosomes for upregulated pexophagy may be limiting the degradation of other substrates.

### Aggrephagy-prone STH*dh^Q111/^*^*111*^ cells have limited pexophagy

If availability of the sequestering autophagosome is the limiting step during upregulated pexophagy, it follows that an increase in any other form of selective autophagy should similarly retard the degradation of other substrates. To test this idea, we next investigated whether the induction of a different form of selective autophagy would have similar effects. Specifically, we asked whether upregulated aggrephagy can influence pexophagy using an aggrephagy-prone cell model of Huntington Disease (HD). HD is an inherited disease caused by an expansion of CAG repeats (>37) in Exon1 of the huntingtin gene, which results in the expression of mutant huntingtin (mHTT) protein with an expanded poly-glutamine tract.^38^ The exact mechanisms by which mHTT expression result in striatal degeneration and HD pathologies are complex, however it has been shown that mHTT forms inclusions that are in part removed by aggrephagy.^38^

One cell model of HD is a neuronal progenitor cell line established from the striatal primordia of knock-in mice expressing mHTT protein containing 111 (STH*dh^Q111/^*^*111*^) glutamine repeats, or HTT protein containing 7 glutamine repeats (STH*dh^Q7/7^*).^39^ STH*dh^Q111/^*^*111*^ cells have been characterized to express mHTT and exhibit heightened autophagy activity compared to STH*dh^Q7/7^* control cells.^39, 40^ We corroborated these findings by monitoring HTT and LC3 by immunofluorescent staining and immunoblot (Fig. S5). We observed depleted HTT and LC3-II protein levels and increased colocalization of HTT and LC3 in STH*dh^Q111/^*^*111*^ cells compared to STH*dh^Q7/7^* cells (Fig. S5). To investigate autophagic flux, we treated cells with BafA1 and observed a greater turnover of HTT and LC3-II in STH*dh^Q111/^*^*111*^ cells compared to STH*dh^Q7/7^* cells (Fig. S5C-H). Collectively, these data demonstrate that STH*dh^Q111/^*^*111*^ cells have upregulated aggrephagy.

To assess whether upregulation of aggrephagy limits the degradation of other selective autophagy substrates, we measured pexophagy activity in STH*dh^Q111/^*^*111*^ and STH*dh^Q7/7^* cells by ectopically expressing PMP34-GFP-UBKo. At basal levels, we observed no difference in peroxisome abundance between STH*dh^Q111/111^* and STH*dh^Q7/7^* cells (Fig. 8A,B). However, expression of the PMP34-GFP-UBKo construct in STH*dh^Q7/7^* cells resulted in a significant decrease in peroxisome number (40% loss of PMP70-labelled peroxisomes), whereas no change was observed in STH*dh^Q111/111^* cells (Fig. 8D-F). These results were supported by immunoblots demonstrating that PMP34-GFP-UBKo expression caused a decrease in PMP70 protein levels in the STH*dh^Q7/7^*, but not the STH*dh^Q111/111^* cells, when compared to cells expressing the non-ubiquitin PMP34-GFP construct (Fig. 8G-I).

**Figure 8.**
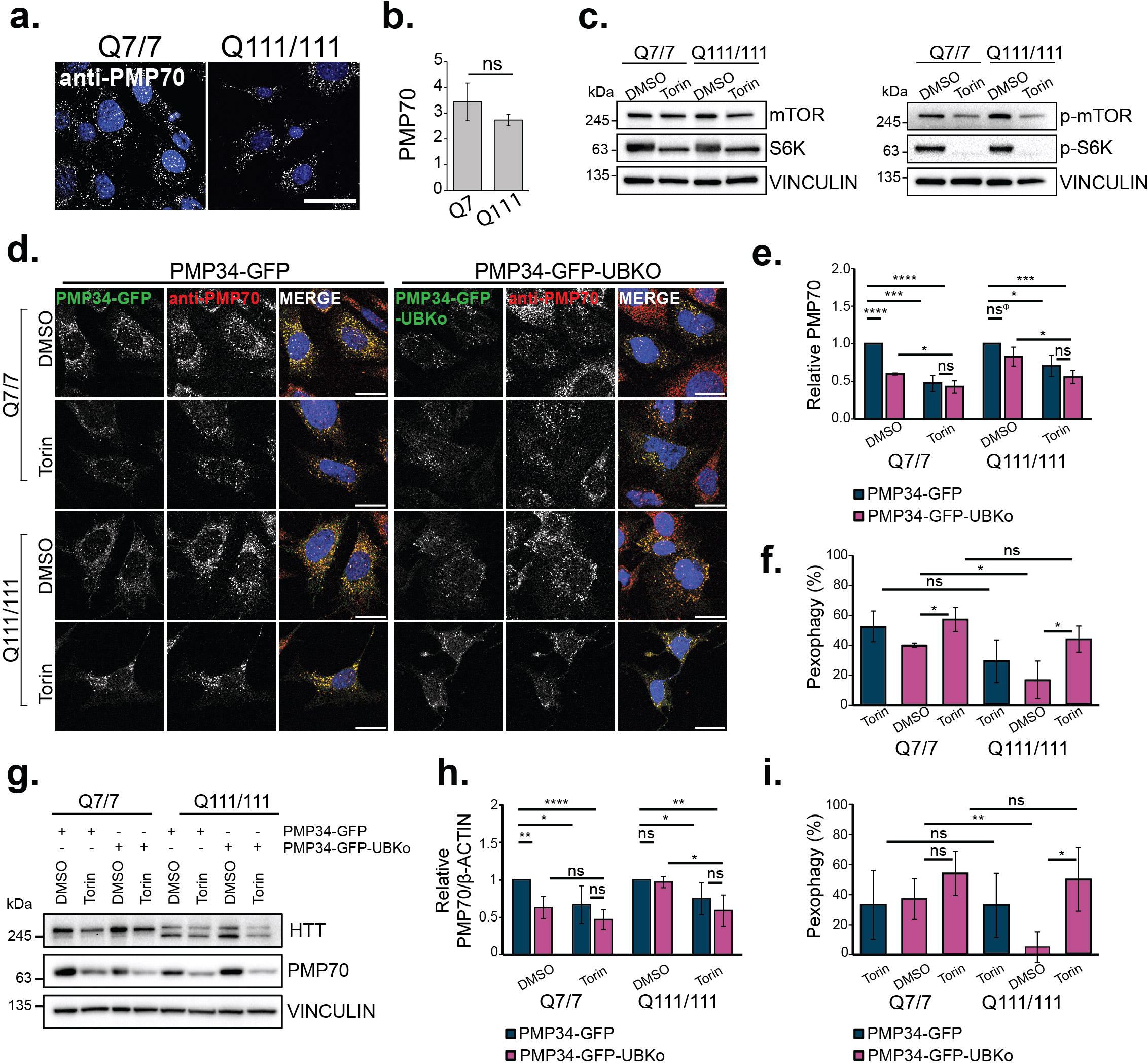
STH*dh^Q111/111^* cells have impaired pexophagy. **(a)** Immunofluorescent images of STH*dh^Q7/7^* or STH*dh^Q111/111^* cells stained for peroxisomal marker, PMP70. **(b)** Quantification of PMP70 in (a). PMP70 was calculated by dividing the number of PMP70 puncta by cell volume. **(c)** Immunoblots of STH*dh^Q7/7^* or STH*dh^Q111/111^* cells treated for 24-h with 100nM Torin1 and probed for mTOR, S6K, p-mTOR, p-S6K, and loading control Vinculin. **(d)** Representative images of STH*dh^Q7/7^* or STH*dh^Q111/111^* cells expressing either PMP34-GFP or PMP34-GFP-UBKo for 48-h, and immunostained for PMP70. Cells were grown in either DMSO or 100nM Torin1 for the final 24-h. **(e)** Quantification of PMP70 in (d), relative to DMSO PMP34-GFP condition for each cell type. **(f)** Quantification of pexophagy in (d). Pexophagy was calculated as the percentage loss of PMP70 from DMSO PMP34-GFP conditions. **(g)** Immunoblots of STH*dh^Q7/7^*or STH*dh^Q111/111^*cells from (a), probed for HTT, PMP70, and loading control Vinculin. **(h)** Densitometry quantifications of PMP70 bands normalized to Vinculin and relative to DMSO PMP34-GFP condition for each cell type. Results represent the mean from n=5 (immunoblot) or n=4 (immunofluorescence) independent trials, and error bars represent standard deviation. For all statistics, two-tailed unpaired student t-test were performed, and significance is indicated by asterisks; *P < 0.05, **P <0.01, ***P<0.001, ****P<0.0001. Scale bars, 25µm.

We next posed whether induction of autophagosomes could improve pexophagy in STH*dh^Q111/111^* cells. PMP34-GFP-UBKo expressing cells were supplemented with Torin1 for 24-h, which robustly inhibited mTORC1 activity (Fig. 8C). Torin1 improved pexophagy activity in STH*dh^Q111/111^* cells compared to DMSO conditions, as shown by increased loss of PMP70 punctate structures in immunofluorescent images and PMP70 protein levels (Fig. 8D-I). Collectively, these data support that upregulation in aggrephagy in the STH*dh^Q111/111^* have decreased capacity for pexophagy compared to STH*dh^Q7/7^* cells, which can be improved by stimulating autophagosome induction.

Complex autophagic dysfunction has been documented in HD models, and wild-type HTT protein has been proposed to mediate the selective autophagy of protein aggregates, lipid droplets, and mitochondria in response to amino acid starvation and other autophagic stimuli.^38, 41–43^ As such, we investigated whether the wild-type HTT protein was required for pexophagy by examining both amino acid starvation and PMP34-GFP-UBKo-induced pexophagy in HeLa cells depleted of HTT. In basal culture conditions, treatment with HTT-targeting siRNA achieved a 93% reduction in HTT protein expression but had no effect on peroxisome abundance (Fig. S6A-F). Following 24-h amino acid starvation, we observed similar loss of peroxisomes in cells treated with mock, non-targeting, or HTT-targeting siRNA, as indicated by a decrease in PMP70 and Catalase protein levels in starved cells compared to basal conditions (Fig. S6A-D). Further, expression of PMP34-GFP-UBKo in cells caused similar peroxisome loss compared to expression of PMP34-GFP in mock, control, or HTT depleted cells (Fig. S6G-I). Conversely, depletion of ATG12 prevented peroxisome loss following expression of PMP34-GFP-UBKo (Fig. S6F-H). Taken together, these findings suggest that the loss of functional HTT protein does not impair pexophagy, and instead supports that increased aggrephagy in STH*dh^Q111/111^* cells influences pexophagy.

## Discussion

A hallmark of neurodegenerative disease is proteinopathy, the accumulation of misfolded proteins that results in cell damage or death. Along with the buildup of pathogenic protein assemblies, neurons also accumulate damaged organelles including mitochondria, lysosomes and ER.^44–46^ Although these pathogenic proteins can themselves induce organelle damage, it is not fully understood why autophagy cannot clear these damaged organelles. In this study, we examined whether upregulation of one selective autophagy pathway can influence the degradation of autophagy substrates. Using pexophagy as a model, we found that upregulated pexophagy limits the autophagic clearance of mitochondria, ALIS, and αS. Similarly, upregulated aggrephagy limits the degradation of peroxisomes. Collectively, our data indicates that an increase in the degradation of a single autophagy substrate can limit the degradative capacity of other selective autophagy pathways.

Our findings suggest that the upregulation of pexophagy exhausts the available sequestering membrane in the cell to limit the degradation of other autophagy substrates. This hypothesis is supported by the ability of rapamycin to improve mitophagy and aggrephagy activity during upregulated pexophagy. Acute rapamycin treatment substantially increased autophagosome flux, suggesting that the rapamycin-rescue of mitophagy and aggrephagy is facilitated by autophagosome induction.

It is likely that selective autophagy is limited downstream of substrate ubiquitin designation. Both proteasomal degradation of OMM proteins as well as Parkin recruitment to mitochondria was observed in PEX1 and PEX13 depleted cells following mitophagy induction, supporting that depolarized mitochondria were ubiquitinated. Moreover, in cells expressing PMP34-GFP-UBKo where aggrephagy was induced, the ubiquitin-binding antibody localized to both PMP34-GFP-UBKo and ALIS, indicating that both substrates were ubiquitinated. Therefore, it is unlikely that substrate ubiquitination is a limiting event during upregulated pexophagy.

Other studies have investigated the rate-limiting steps(s) of autophagy. An investigation of ROS-induced mitophagy found that the rate-limiting step of mitochondria degradation was delayed lysosomal fusion or acidification.^47^ Additionally, a study of autophagy rates found that rapamycin treatment results in a robust increase in the rate of autophagosome formation but a lag in rate of autolysosome fusion and degradation.^48^ In our study, we observed that lysosome acidification was in fact increased during upregulated pexophagy, and found no impairments in autophagosome flux following Baf A1 treatment suggesting that autolysosome fusion is not affected. Further, only a prolonged 24-h rapamycin treatment, and not a 5-h treatment (data not shown) increased cellular lysosomes, supporting that the rapamycin rescue of mitophagy and aggrephagy in our cell assays was not mediated by increased lysosomes. Collectively, our findings suggest that lysosome availability is not limiting during upregulated pexophagy.

The idea of competition during selective autophagy has yet to be addressed in the field. During 36-h amino acid starvation, it was shown that there appears to be an ordered degradation of autophagy substrates, beginning with degradation of cytosolic and proteasomal proteins, then ribosomes, then finally large organelles including the ER and mitochondria.^49^ Whether certain autophagy substrates are prioritized for degradation during other autophagic stimuli has yet to be elucidated. From our study, it appears that the priority for substrate degradation is related to the temporal induction of each selective autophagy pathway. As pexophagy was induced for 24-48h prior to pharmacological induction of mitophagy or aggrephagy in our assays, it can be speculated that at the time of mitophagy or aggrephagy induction, peroxisomes have sequestered much of the autophagy machinery and are thus degraded first.

Notably, a previous study by Lee *et al*. reported that the loss of PEX13 results in impaired autophagic clearance of mitochondria and Sindbis virus.^21^ The defects in mitophagy and virophagy were attributed to PEX13 having a direct role in these processes. However, we have recently demonstrated that PEX13 is required to prevent pexophagy of healthy peroxisomes, and that it is degraded to mediate pexophagy during starvation.^17^ Here, we show that PEX13 alone is not responsible for preventing mitophagy or aggrephagy, as artificially targeting a ubiquitin motif to peroxisomes to induce pexophagy was sufficient to retard aggrephagy. A potential explanation as to why Lee *et al.* may have missed the role of PEX13 in pexophagy is the lack of detailed analysis of changes in peroxisomes.

Our work in ZSD PEX1^G843D^ patient fibroblasts shows that they exhibit impaired aggrephagy that can be rescued with pharmacological induction of autophagy. Combined with our evidence that increased pexophagy impairs mitophagy and αS clearance, we propose that selective autophagy may be limited in ZSD. This hypothesis is supported by existing ZSD mouse models with dysregulated pexophagy that yield an accumulation of autophagy substrates, such as damaged mitochondria and αS. For example, Rahim, R.S. *et al*. report a five-fold increase in mitochondrial volume with increased oxidative stress and impaired function in a brain-restricted PEX13 knockout mouse model.^14^ While this expansion of the mitochondrial compartment may in part be explained by the reported increase in PGC-1α-mediated mitochondrial biogenesis, the accumulation of mitochondria with impaired function in particular suggests that autophagy is not effectively removing damaged portions of mitochondria.^14^ Additional ZSD mouse models with dysregulated pexophagy demonstrate an increase in the oligomerization and phosphorylation of αS, events that precede Lewy Body formation in PD and should be alleviated by selective autophagy.^13, 27^

Taken together, we propose that in ZSD cases arising from increased pexophagy, the loss of peroxisomes has multifactorial consequences on cellular health. First, the loss of peroxisome function bears a host of mitochondrial insults including peroxin mislocalization, increased oxidative stress, in addition to a reduction in fatty acid oxidation and respiration rates.^13–15, 50, 51^ Alongside, the sustained increase in pexophagy limits selective autophagy, and as a result populations of damaged mitochondria and αS oligomers accumulate. It follows that pharmacologically stimulating autophagy may improve the health of the mitochondrial network in ZSD patients.

As this study focused on how upregulated pexophagy influences selective autophagy, our conclusions are largely constrained to conditions of increased pexophagy. However, we predict that saturation of selective autophagy is not limited to pexophagy but the upregulation of other selective autophagy pathways. Our work in the HD cell model bares some support for this hypothesis. However, although our studies show that HTT is not required for pexophagy, it remains a possibility that mHTT affects pexophagy through mechanisms other than depleting the available autophagosomes. Whether an increase in the degradation of different selective autophagy substrates, such as mitochondria, ALIS, or ribosomes, can similarly limit selective autophagy awaits further investigation. Determining how the cell coordinates concurrent selective autophagy pathways will provide insights into the pathogenesis of ZSD and other diseases that are associated with selective autophagy impairments. One example is severe acute malnutrition, where hepatic dysfunction in a rat model of childhood malnutrition was characterized by the loss of peroxisomes via increased pexophagy, followed by accumulation of dysmorphic and dysfunctional mitochondria.^52^ In this study of severe childhood malnutrition, we hypothesize that increased pexophagy may limit selective autophagy and hinder the clearance of damaged mitochondria. More broadly, neurodegenerative proteinopathies burdened by a chronic increase in aggrephagy may experience limited selective autophagy. Although complex mechanisms by which misfolded proteins such as αS and mHTT contribute to organelle and autophagy dysfunction have been described, we hypothesize that limited selective autophagy may further hinder the removal of damaged components.^53, 54^

Crosstalk between degradative pathways helps to protect cells from proteotoxic and metabolic stress. For example, inhibition of the proteasome in cultured neural cells or chaperone-mediated autophagy (CMA) in mouse fibroblasts or liver results in a compensatory upregulation of macroautophagy.^55–57^ The secretion of autophagy substrates upon defective autophagy has also been recently described, and could be a mechanism by which limited selective autophagy may exacerbate tissue damage, if damaged components targeted to the lysosome are instead secreted into the extracellular space.^58, 59^ How these degradative pathways influence one another across tissues and metabolic states remains a key area to address.

Numerous selective autophagy pathways including aggrephagy, mitophagy, pexophagy, reticulophagy, ribophagy, and lipophagy have been characterized in the last decade, generating a new appreciation for the selectivity of autophagy processes. Here, we have provided evidence to support that these selective autophagy pathways can influence each other, and that the degradative capacity of selective autophagy can be limited in neurodegenerative disease models. In future studies, we hope to address how the cell regulates selective autophagy pathways in parallel to ensure the proper degradation of autophagy substrates using *in vitro* and *in vivo* models.

## Materials and Methods

The reagents used in this study including chemicals, antibodies, and commercially available kits are summarized in Supplementary Table 1.

### Constructs and siRNA

The PMP34-GFP, PMP34-GFP-UBko and LC3-Cherry constructs used in this study were previously described.^23, 26^ LAMP1A-mEmerald was donated by Dr. Sergio Grinstein (Hospital for Sick Children, Toronto, ON, Canada). The siRNAs used in this study were custom synthesized from Sigma-Aldrich: PEX1 (5’-CCAAG CAACUUCAGUCAAA-3’), PEX2 (5’-CUCUUACUGGUGCACCUAA-3’), PEX13 (5’-CGGUAUCUUUACAGACGGCUAC-3’), PEX14 (5’-GAACUCAAGUCCGAAAUU-3’), ATG12 (5’-GUGGGCAGUAGAGCGAACA-3’), NBR1 (5’-GGAGUGGAUUUACCAGUUAUU-3’), HTT (AACUUUCAGCUACCAAGAAAG) and non-targeting control (5’- AAUAAGGCUAUGAAGAGAUAC-3’). siRNA knockdown was validated by immunoblot or TaqMan real-time quantitative PCR.

### Cell culture and Transfection

HeLa cells were purchased from American Type Culture Collection (CCL-2). Stably transfected GFP-Parkin HEK293 cells were a gift from Dr. Angus McQuibban (University of Toronto, Toronto, ON, Canada), and are previously described.^60^ The immortalized patient-derived skin fibroblast cell lines from healthy (Control), PEX1^G843D^, PEX3-R53ter, and PEX19-deficient (PBD399-T1) patients were gifts from Dr. Nancy Braverman (McGill University, Montreal, QC, Canada) and S.J Gould (John Hopkins University School of Medicine, Baltimore, MD). STH*dh^Q111/111^* and STH*dh^Q7/7^* cells were donated by Dr. Ray Truant (McMaster University, Hamilton, ON, Canada), and are previously described.^39^ All cells were cultured in Dulbecco’s modified Eagle’s medium (Gibco) supplemented with 10% fetal bovine serum (Wisent), at 5% CO_2_ in a 37°C humidified incubator. Cells were routinely tested for mycoplasma (FroggaBio, 25235). For amino acid starvation experiments, cells were washed twice with PBS and subjected to HBSS for the indicated time (Lonza). siRNA and plasmid transfections were performed using Lipofectamine 2000 (Invitrogen) according to the manufacturer’s instructions. For all knockdown experiments, a 2-day siRNA transfection was performed followed by a 24-h recovery period before chemical treatments were applied. In experiments where cells were transfected with plasmids, plasmids were expressed for 24-48h preceding chemical treatments, as specified in the figure captions. For experiments performed in STH*dh^Q111/111^* and STH*dh^Q7/7^* cells as well as the lysosome experiments performed in HeLa cells, transfection was performed using the Neon transfection system (Invitrogen) according to the manufacturer’s instructions.

### Immunoblotting

Cells were washed with PBS and lysed in 100mM Tris, pH 9, 1% SDS with 1X protease cocktail inhibitor (BioShop, PIC002.1). For analysis of phosphorylated proteins, lysis buffer was supplemented with 20mM NaF and 5mM Na_2_VO_4_. Lysates were boiled for 15-min at 95°C with vortexing every 5-min, and then centrifuged at 13,000 *g* for 15-min. Protein concentration of the supernatant was measured and equivalent sample amounts were subjected to SDS-PAGE. Protein was transferred to a PVDF membrane and blocked for 1-h in 5% skim milk. Membranes were incubated in the appropriate primary and secondary antibodies diluted in 1% skim milk or 3% BSA. Proteins were visualized using Amersham ECL Prime Western Blot Detection Reagent (VWR, CA89168-782) and a ChemiDoc Imaging System (Bio-Rad Laboratories).

### Immunofluorescence

Cells were seeded on no.1 glass coverslips (VWR, CA-89015725). For fixation, cells were washed with PBS and incubated in 3.7% paraformaldehyde (Electron Microscopy Sciences) for 15-min. When indicated, cells were treated with 0.025% digitonin for 10 seconds prior to fixation. Cells were washed twice with PBS and permeabilized with 0.1% Triton X-100 for 15-min. After two final PBS washes, cells were incubated in blocking buffer (10% FBS in PBS) for 1-h at room temperature. For mitochondrial staining, cells were fixed in warm 3.7% paraformaldehyde containing 5% sucrose. Cells were then incubated with the appropriate primary and secondary antibodies diluted in blocking buffer at room temperature for 1-h each. For DAPI staining, cells were incubated in DAPI diluted 1:10,000 in PBS for 15-min. Coverslips were mounted on glass slides using DAKO mounting medium and stored at 4°C. Fluorescence microscopy was performed on either a Zeiss LSM710 or LSM980 laser-scanning confocal microscope with a 63x/1.4 NA Plan-APOCRAMAT oil immersion objective. Images were taken using a combination of 6 laser lines (405, 458, 488, 514, 561, 633 nm) and the appropriate filters. *Z*-stacks were acquired using ZEN 2009 or ZEN 2019 (Zeiss Enhanced Navigation) software.

For each experimental trial, all images were acquired using the same acquisition setting with minimal saturated pixel. Images used in figures were selected as representative images which best reflect the quantified data and were adjusted for brightness/contrast for presentation purposes only using Adobe Photoshop CS6.

### Quantification and Statistical Analyses

Immunofluorescence quantification analyses were conducted using Volocity software (Perkin Elmer). Quantifications were generated by drawing a region of interest (ROI) around individual cells and applying the ‘Find Objects’ algorithm in Volocity, using thresholding to control for size and intensity cut-off. Cell volume was recorded from the ROI. For peroxisome quantification, individual PMP70 puncta were counted and the number of PMP70 structures in a given ROI was then divided by cell volume. For ALIS quantification, FK1- or FK2-aggregates in a given ROI were identified based on applied thresholds, and the total volume of FK1- or FK2- aggregates was divided by cell volume to obtain relative aggregate volume. For mitochondria quantification, total fluorescence intensity of CVα was recorded and divided by cell volume. For alpha-Synuclein quantification, total fluorescence intensity of αS was recorded and divided by cell volume. For lysosome quantification, LAMP1A or cresyl violet structures in a given ROI were identified based on applied thresholding and the total volume of LAMP1A or cresyl violet structures was divided by cell volume. For co-localization analyses, Pearson’s Correlation coefficients were recorded from Volocity. For each experiment, at least three separate trials (n=3) were conducted, and average values were calculated by averaging the appropriate densities of at least 25 cells per trial.

Immunoblot densitometry quantifications were performed using ImageJ software, and presented as an average of at least three (n=3) independent trials. Each protein band is normalized to its relevant probed loading control, either GAPDH, β-Actin, or Vinculin.

Statistical analyses were performed using either Student’s *t*-test or One-Way ANOVA where appropriate using and is indicated in each figure caption. Statistical tests were performed using either Microsoft Excel or GraphPad Prism 5 to include all available data for each experiment. The distribution of the data were assumed normal.

### Mitophagy Assay

Mitophagy was induced in GFP-Parkin HEK293 for 8-h with 2.5µm oligomycin and 250nM antimycin A1, and 2µM rapamycin or 200nM bafilomycin A1 where applicable. Cells were either lysed for immunoblotting, or fixed and immunostained for CVα, and analysed with confocal fluorescent microscopy. Mitochondrial clearance was assessed by quantifying CVα relative fluorescence intensity, or calculating the percentage loss of mitochondrial proteins from densitometry quantifications of immunoblots.

### Aggrephagy Assay

Cells were treated for 3-h with 5µg mL^-1^ puromycin to induce ALIS. Cells were washed with PBS and allowed a 5-h clearance period in DMEM, or 2µM rapamycin where applicable. Cells were fixed and immunostained for FK1 or FK2, and analysed with confocal fluorescent microscopy. Aggrephagy was assessed by quantifying relative FK1/FK2-ALIS volume.

### Alpha-Synuclein Clearance Assay

αS PFF were generated using recombinant human αS expressing the A53T mutation associated with familial Parkinson’s Disease. Briefly, 1mg mL^-1^ αS was incubated for 7 days at 37°C with constant agitation in a buffer of 20mM Tris-HCl, pH 7.4 and 100mM NaCl, as previously described.^62^ Fibril formation was confirmed with a ThT fluorescence assay. For the αS clearance assay, αS PFF were transfected for 4-h at a final concentration of 1µg mL^-1^ using Lipofectamine 2000, according to the manufacturer’s protocol. Cells were washed with PBS and allowed an 8-h clearance period in DMEM, 2µM rapamycin, or 0.25mM Leupeptin and 2µM E-64 where applicable. Cells were fixed and immunostained for αS, and analysed with confocal fluorescent microscopy. αS clearance was assessed by quantifying αS relative fluorescent intensity in cells.

### Lysosome Assay

HeLa cells were transfected with siRNA and DNA together by electroporation, seeded in LabTek^TM^ chambers (Nunc) and grown for 48-h. Prior to imaging, cells were washed twice, incubated for 5-min in 1.5µM cresyl violet (Sigma-Aldrich) in HBSS, and washed twice again. Cells were imaged live in a temperature (37`°`C) and atmosphere (5% CO_2_) controlled environment, and all images were acquired within 20-min of cresyl violet treatment. Cells were treated with rapamycin or torin1 for 24-h (100nM) or 5-h (2µM) where applicable. To dissipate lysosomal acidification, cells were pretreated for 1-min with 30mM NH_4_Cl and 250nM CcA in HBSS and then incubated for 5-min with 1.5µM cresyl violet in the same solution.

### Pexophagy Assay

PMP34-GFP or PMP34-GFP-UBKo was expressed in cells for 48-h prior to fixation. For torin1 experiments, cells were supplemented with 100nM torin1 for the final 24-h. Cells were either fixed and immunostained for PMP70 and analysed with confocal microscopy, or lysed for immunoblotting. Pexophagy was assessed by quantifying the percentage loss of PMP70 from PMP34-GFP to PMP34-GFP-UBKo conditions.

## Acknowledgments

This work was supported by the Canada Institute of Health Research grants to P.K.K (PJT #156196) and to R.B (PJT# 156307), as well as the Ontario Graduate Scholarship, Hayden Hantho Award, and the Hilda and William Courtney Clayton Paediatric Research Fund to K.G.

## Author Contributions

All authors contributed to the data analysis and interpretation of the data. K.G., R.B, and P.K.K. conceived the study. K.G., J.C.W., R.B, and P.K.K. designed the experiments. K.G. performed the experiments, R.W.L. generated the αS fibrils. K.G, R.B. & P.K.K wrote the manuscript. All authors contributed to editing of the manuscript.

## Ethics declarations

The authors declare no competing interests.

## Extended Data Figure Legends

**Extended Data Figure 1.**
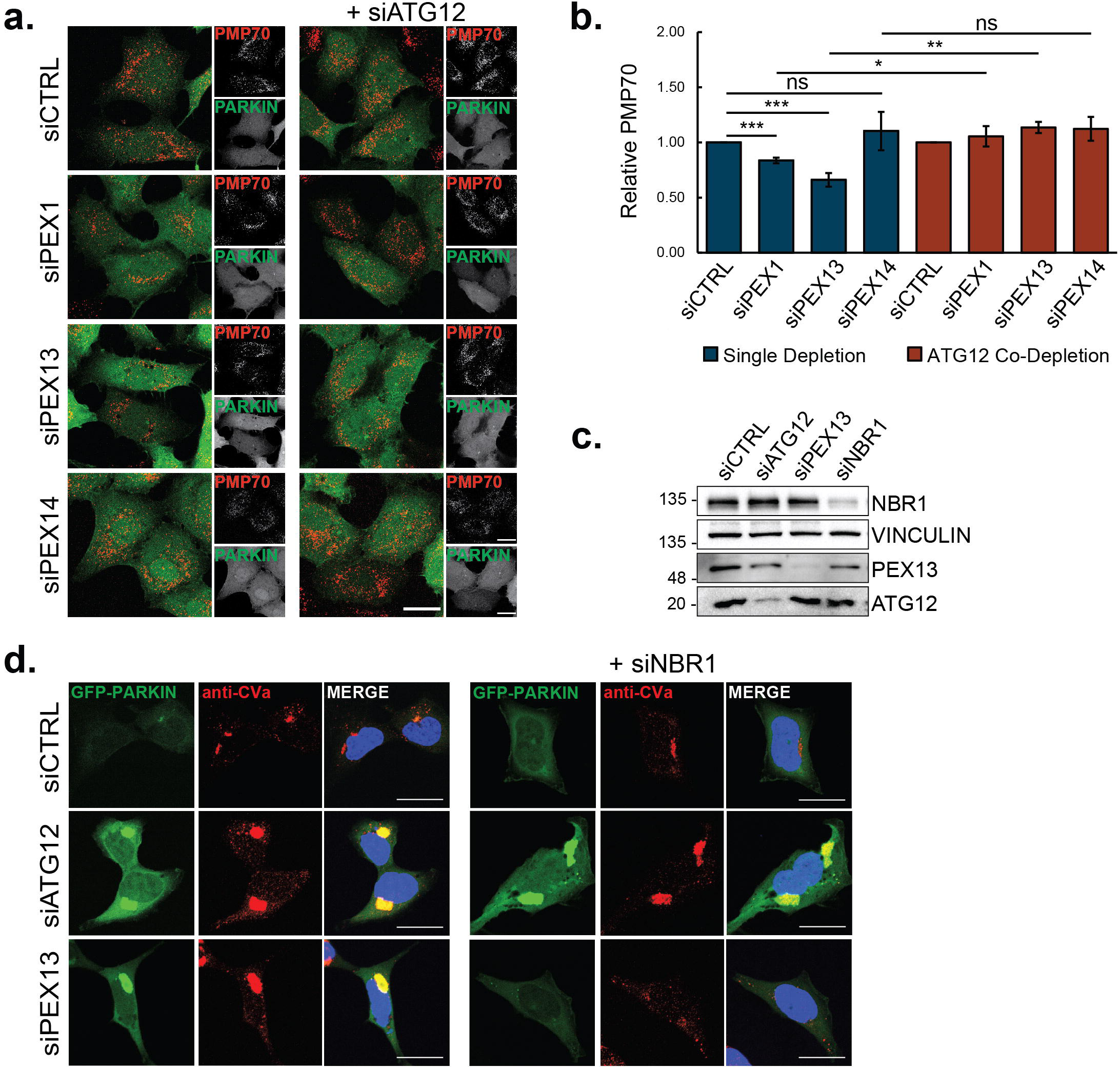
PEX1 or PEX13 depletion increases pexophagy and impairs mitophagy in GFP-PARKIN HEK cells. **(a)** Immunofluorescent images of GFP-PARKIN HEK cells treated with the indicated siRNA and immunostained for peroxisomal marker, PMP70. Scale bar, 25µm. **(b)** Quantification of PMP70 in (a), relative to control siRNA treated cells. PMP70 was measured by dividing the number of PMP70 puncta by cell volume. Results represent mean from *n*=3 independent trials (30 cells quantified/trial) and error bars represent standard deviation. *P < 0.05, **P <0.01; two-tailed unpaired student t-test. **(c)** Immunoblot of GFP-PARKIN HEK cells treated with the indicated siRNA and probed for NBR1, PEX13, and ATG12. **(d)** Representative images of GFP-Parkin HEK cells treated with the indicated siRNA(s) prior to treatment for 8-h with 2.5µM Oligomycin and 250nM Antimycin A1. Cells were immunostained for mitochondria marker, CVα. Scale bars, 25µm.

**Extended Data Figure 2.**
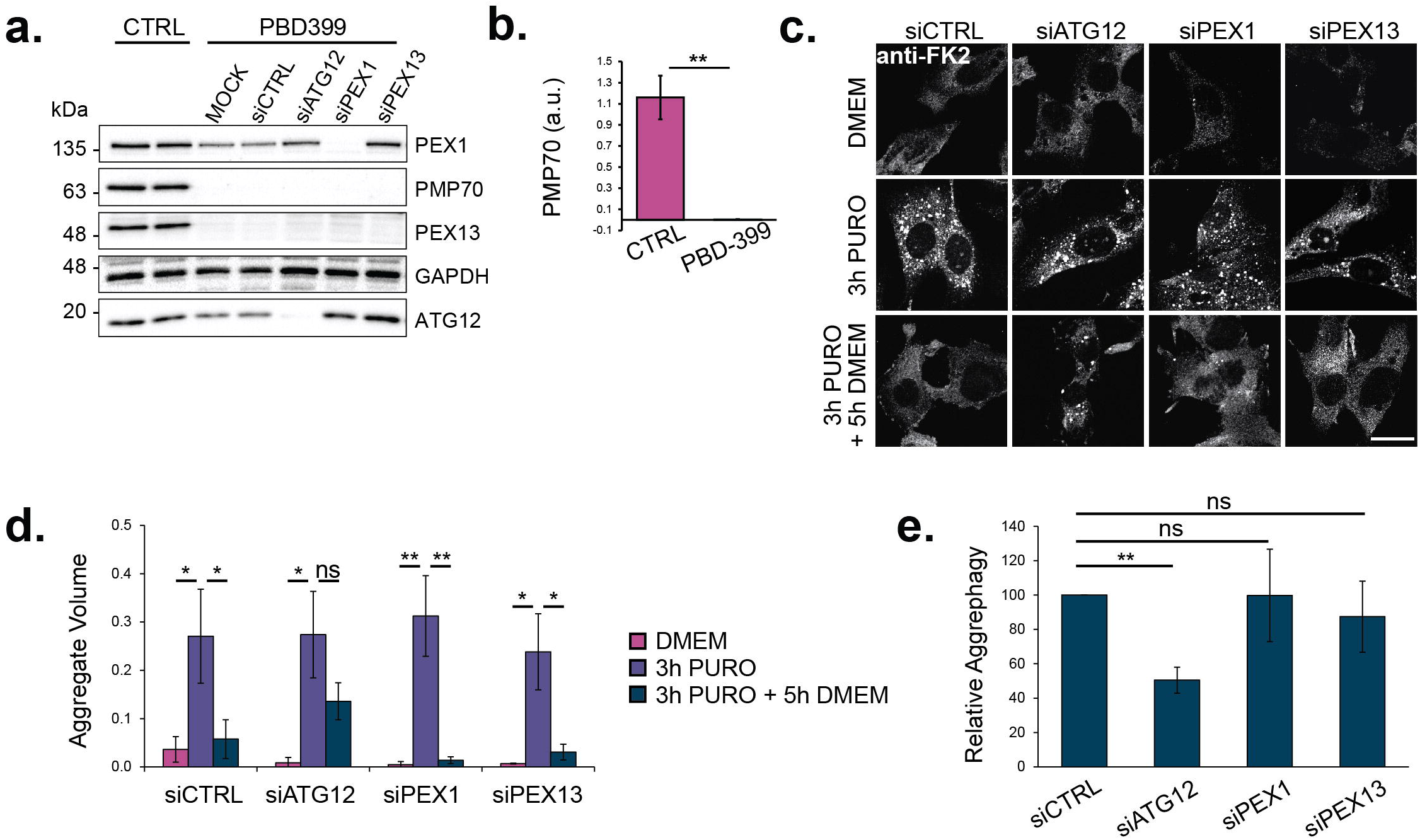
Peroxisome-deficient fibroblasts have unimpaired aggrephagy. **(a)** Immunoblot of Control and PBD399-T1 fibroblast cells grown treated with siRNA and probed for the indicated proteins. **(b)** Densitometry quantification of PMP70 bands normalized to loading control GAPDH. Bands were quantified using ImageJ software from *n=3* independent trials. **(c)** PBD399-T1 fibroblasts treated with the indicated siRNA prior to aggrephagy induction. Representative images from cells at each stage in the assay: DMEM, 3-h 5µg mL^-1^ Puromycin, 3-h 5µg mL^-1^ Puromycin followed by 5-h clearance period in DMEM. Cells were immunostained for FK2. Scale bars, 25µm. **(d)** Quantification of Relative Aggregate Volume in (c). Relative Aggregate Volume was calculated by dividing the total volume of FK2 puncta by cell volume. **(e)** Quantification of aggrephagy activity in (c), relative to siRNA-control-transfected cells. Aggrephagy activity was measured as the % clearance of FK2 aggregates. Results represent mean from *n*=3 independent trials (30 cells quantified/trial) and error bars represent standard deviation. *P < 0.05, **P <0.01, ***P<0.001; two-tailed unpaired student t-test.

**Extended Data Figure 3.**
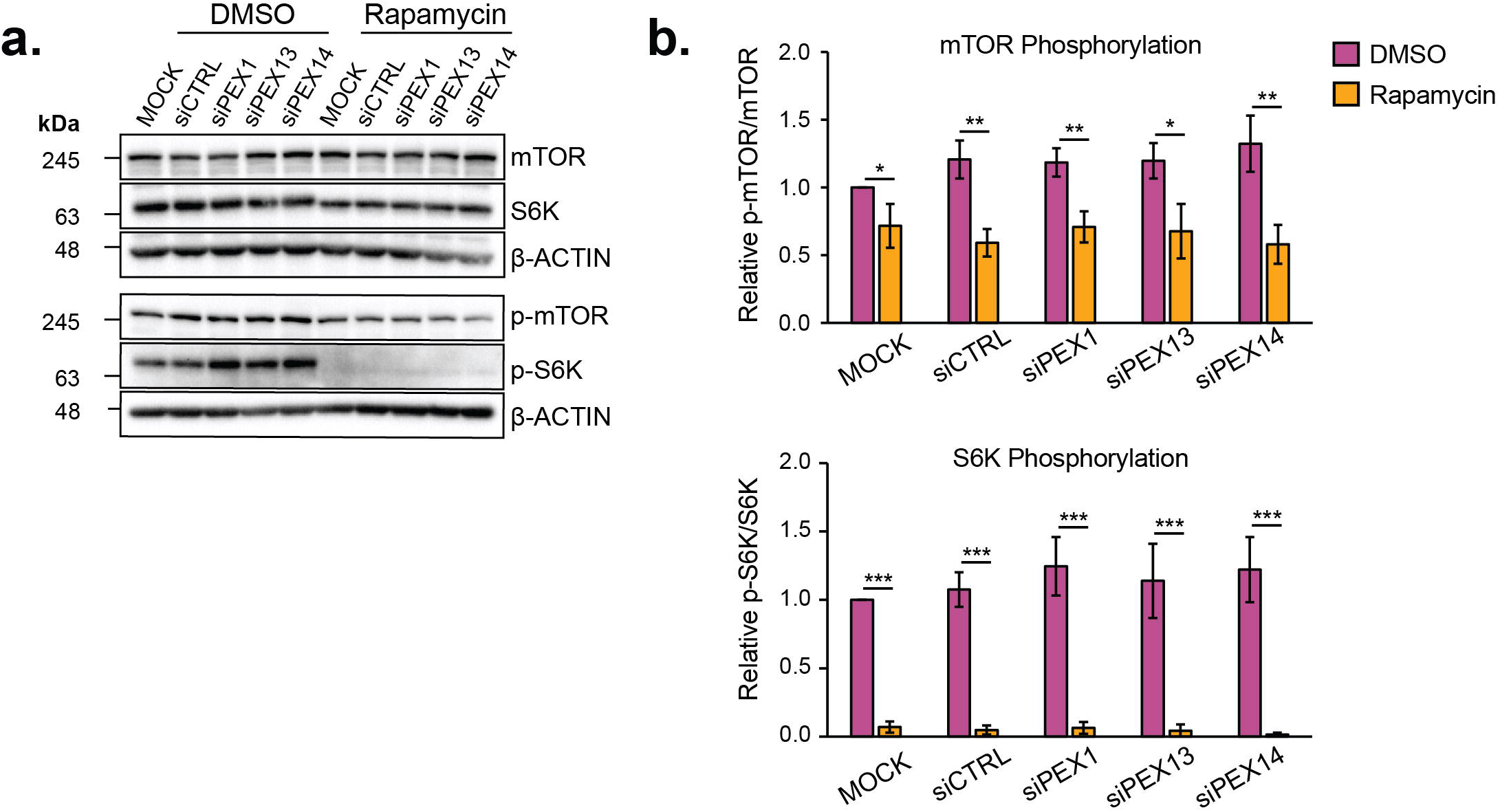
Rapamycin treatment inhibits mTOR in PEX1 and PEX13 depleted cells. **(a)** Immunoblots of HeLa cells treated with the indicated siRNA and grown in either DMSO or 2µM Rapamycin for 5-h. Immunoblots were probed for mTOR, SK6, phospho-mTOR, phospho-SK6, and loading control β-Actin. Quantification of **(b)** mTOR and **(c)** S6K phosphorylation from (a), relative to MOCK. mTOR and S6K phosphorylation were calculated by dividing the band intensity of phospho-mTOR/mTOR or phospho-SK6/S6K. Band intensities were first measured using ImageJ software and normalized to their respective β-Actin loading control. Results represent mean from *n*=3 independent trials and error bars represent standard deviation. *P < 0.05, **P <0.01, ***P<0.001; two-tailed unpaired student t-test.

**Extended Data Figure 4.**
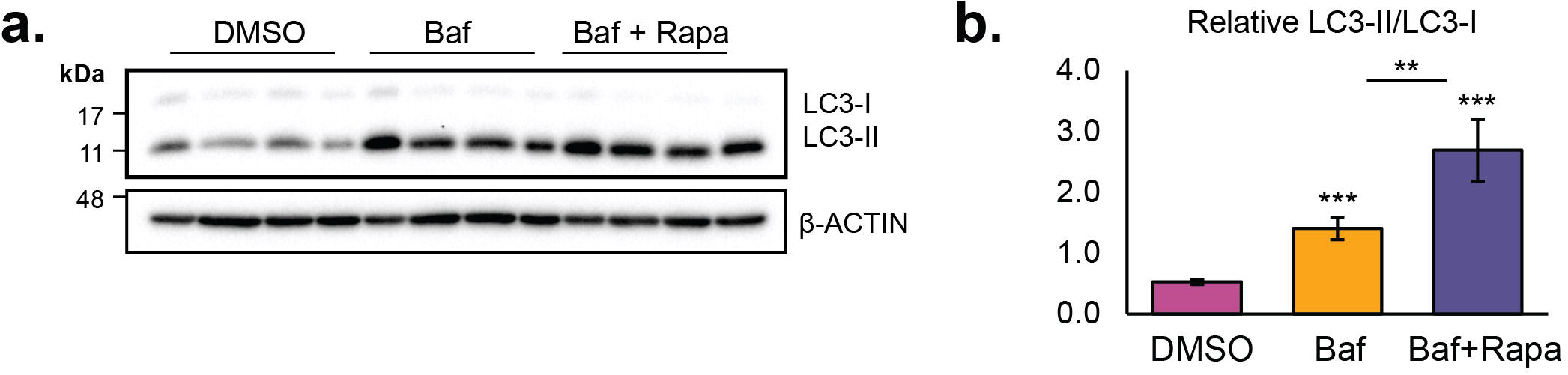
Rapamycin treatment induces autophagy flux in PEX14 depleted cells. **(a)** Immunoblot of HeLa cells treated with siRNA targeting PEX14 and grown in either DMSO, 200nm Bafilomycin, or 2µM Rapamycin + 200nM Bafilomycin A1 for 5-h. 4 lanes per condition in the immunoblot correspond to samples from *n*=4 independent trials. **(b)** The mean densitometry quantification of LC3-II/LC3-I ratio in (a), relative to MOCK DMSO condition. Error bars represent standard deviation. **P <0.01, ***P<0.001; two-tailed unpaired student t-test.

**Extended Data Figure 5.**
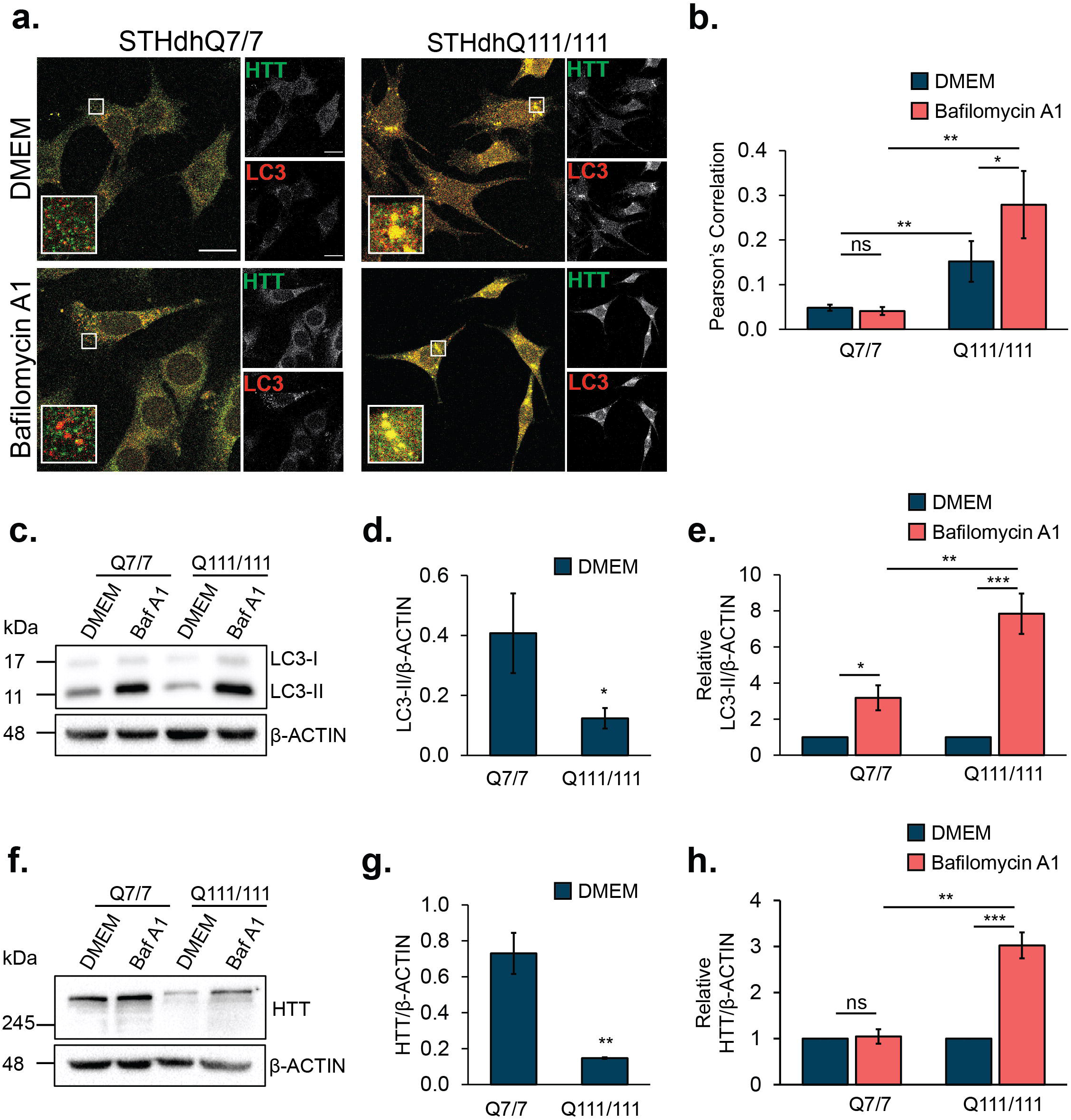
STH*dh^Q111/111^* cells undergo increased Huntingtin turnover by autophagy. **(a)** Immunofluorescent images of STH*dh^Q7/7^* or STH*dh^Q111/111^* cells grown in DMEM or 0.25mM Bafilomycin A1 for 24-h and immunostained for Huntingtin (HTT) and LC3. Scale bar, 25µm. White squares denote zoomed region. **(b)** Pearson’s correlation coefficient for LC3 and HTT from (a). **(c)** Immunoblot of STH*dh^Q7/7^*or STH*dh^Q111/111^* cells grown in DMEM or 0.25mM Bafilomycin A1 for 24-h and probed for **(c)** LC3 or **(f)** HTT. Quantification of **(d)** LC3-II or **(g)** HTT bands normalized to β-Actin for DMEM condition. All immunoblot bands were quantified using ImageJ software from *n=3* independent trials. Quantification of normalized **(e)** LC3-II or **(h)** HTT bands relative to DMEM condition for each cell type. Results represent mean from *n*=3 independent trials (30 cells quantified/trial) and error bars represent standard deviation. *P < 0.05, **P <0.01, ***P<0.001; two-tailed unpaired student t-test.

**Extended Data Figure 6.**
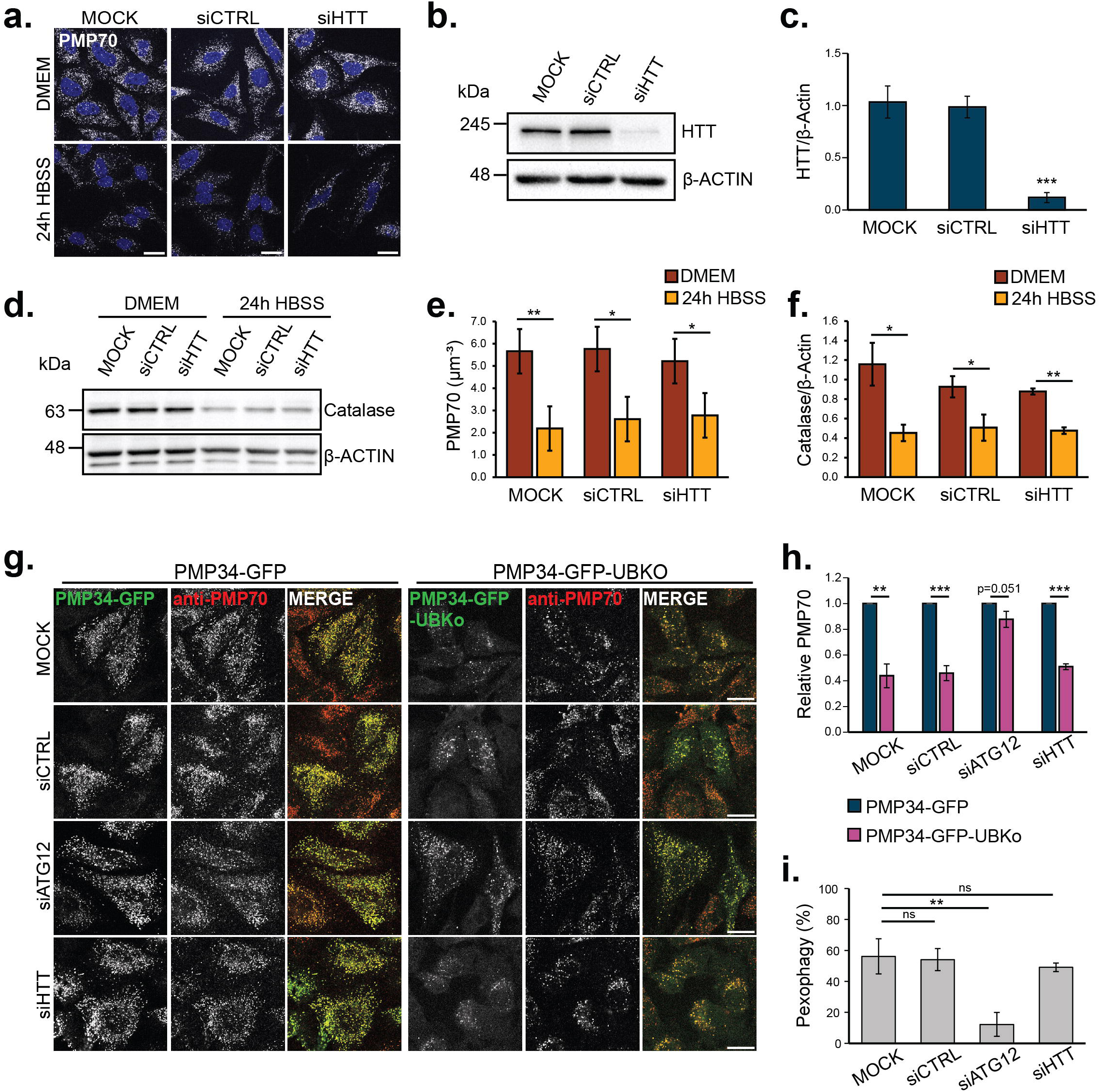
HTT depletion does not influence pexophagy. **(a)** Immunofluorescent images of HeLa cells treated with the indicated siRNA and grown in either DMEM or HBSS for 24-h. Cells are immunostained for PMP70 and nuclei are visualized with DAPI. **(b)** Immunoblot of HeLa cells treated with the indicated siRNA and probed for HTT and β-Actin. **(c)** Quantification of HTT bands normalized to β-Actin. **(d)** Immunoblot of HeLa cells treated with the indicated siRNA and grown in either DMEM or HBSS for 24-h, and probed for Catalase. **(e)** Quantification of PMP70 in (a). PMP70 was measured by dividing the number of PMP70 puncta by cell volume. **(f)** Quantification of Catalase bands normalized to β-Actin. **(g)** Immunofluorescent images of HeLa cells treated with the indicated siRNA and transiently expressing either PMP34-GFP or PMP34-GFP-UBKo for 48-h. Cells are immunostained for PMP70. **(h)** Quantification of PMP70 in **(g)**, relative to PMP34-GFP expressing cells for each siRNA condition. **(i)** Quantification of pexophagy in (g). Pexophagy was quantified as percentage loss of PMP70 with PMP34-GFP-UBKo expression compared to PMP34-GFP. Quantifications represent mean from *n*=3-4 independent trials (30 cells quantified/trial for immunofluorescence). Scale bars, 25µm. Error bars represent standard deviation. *P < 0.05, **P <0.01, ***P<0.001; two-tailed unpaired student t-test.

