## Supplemental Table 1 for "Upregulated pexophagy limits the capacity of selective autophagy"

Supplementary Information: Table 1

| Reagent | Commercial Supplier |
|---|---|
| KITS | |
| SV Total RNA Isolation Program | Promega |
| High-Capacity cDNA Reverse Transcription | Applied Biosystems |
| TaqMan Fast Advanced Master Mix | Applied Biosystems |
| Pierce$^{TM}$ BCA Protein Assay Kit | Thermo Fisher Scientific |
| CHEMICALS | |
| Oligomycin | Fisher Scientific; AAJ61898MA |
| Antimycin A1 | Santa Cruz; sc-202467 |
| Puromycin | BioShop; PUR333 |
| Rapamycin | BioShop; RAP004 |
| Bafilomycin A1 | Santa Cruz; sc-201550A |
| Torin1 | Abcam; ab218606 |
| Leupeptin | BioShop; LEU001 |
| E-64 | Millipore Sigma; E3132 |
| Cresyl Violet Acetate | Sigma-Aldrich; C5042 |
| Concanamycin A | Abcam; ab144227 |
| PRIMARY ANTIBODIES | |
| PMP70 | Abcam; ab3421 |
| PEX1 | BD Biosciences; 611719 |
| PEX13 | Abcam; ab96841 |
| PEX14 | Proteintech; 10594-1-AP |
| ATG12 | Cell Signaling; 2010 |
| NBR1 | Abnova; H00004077-MOI |
| MAP1LC3B | Thermo Fisher Scientific; A-21070 |
| SQSTM1 | BD Biosciences; 610832 |
| CVα | Abcam; ab14748 |
| MFN2 | Abcam; ab56889 |
| HSP60 | Abcam; ab46798 |
| FK2 | Enzo; BML-PW8810 |
| FK1 | Millipore Sigma; 04-262 |
| mTOR | Cell Signaling; 2983 |
| phos-mTOR | Cell Signaling; 2971 |
| p70 S6K | Cell Signaling; 2708 |
| phos-p70 S6K | Cell Signaling; 9205 |
| α-Synuclein | BD Biosciences; 610786 |
| Catalase | Millipore Sigma; 219010 |
| HTT | Millipore Sigma; MAB2166 |
| SECONDARY ANTIBODIES | |
| GAPDH-HRP | Novus Biologicals; NB300-328H |
| β-Actin-HRP | Cell Signaling; 5125 |
| Vinculin-HRP | Cell Signaling; 18799 |
| anti-rabbit IgG-HRP | Thermo Scientific; 31460 |
| anti-mouse IgG-HRP | Cedarlane; CLCC30007 |

| | |
|---|---|
| Anti-mouse IgG Alexa Fluor 488 | Thermo Fisher Scientific; A11001 |
| anti-rabbit IgG Alexa Fluor 568 | Thermo Fisher Scientific; A-11011 |
| anti-mouse IgG Alexa Fluor 568 | Thermo Fisher Scientific; A-11004 |
| anti-mouse IgG Alexa Fluor 647 | Thermo Fisher Scientific; A-31571 |
